# Adjusting for genetic confounders in transcriptome-wide association studies leads to reliable detection of causal genes

**DOI:** 10.1101/2022.09.27.509700

**Authors:** Siming Zhao, Wesley Crouse, Sheng Qian, Kaixuan Luo, Matthew Stephens, Xin He

## Abstract

Expression Quantitative Trait Loci (eQTLs), provide valuable information on the effects of genetic variants. Many methods have been developed to leverage eQTLs to nominate candidate genes of complex traits, including colocalization analysis, transcriptome-wide association studies (TWAS), and Mendelian Randomization (MR)-based methods. All these methods, however, suffer from a key problem: when using the eQTLs of a gene to assess its role in a trait, nearby variants and nearby genetic components of expression of other genes can be correlated with the eQTLs of the test gene, while affecting the trait directly. These “genetic confounders” often lead to false discoveries. We introduced a novel statistical framework to address this challenge. Our method, causal-TWAS (cTWAS), borrowed ideas from statistical fine-mapping, and allowed us to adjust all genetic confounders. In our simulations, we found that existing methods based on TWAS, colocalization or MR all suffered from high false positive rates, often greater than 50%. In contrast, cTWAS showed calibrated false positive rates while maintaining power. Application of cTWAS on several common traits highlighted the weakness of existing methods and discovered novel candidate genes. In conclusion, cTWAS is a novel statistical framework to integrate eQTL and GWAS data, enabling reliable gene discoveries.

## Introduction

Genome-wide association studies (GWAS) have successfully identified a large number of loci associated with a range of complex human traits [1] [2]. It remains difficult, however, to translate these associations into knowledge of causal variants, their target genes and molecular mechanisms [3]. A common strategy to address this challenge is the use of expression quantitative trait loci (eQTL) data to link variants with gene expression. Transcriptome-wide association studies (TWAS) provides a formal framework for the joint analysis of eQTL and GWAS data [4] [5]. In TWAS, researchers build predictive models of gene expression from genetic variants, usually in *cis*, and then test for associations between predicted (“imputed”) expression and traits of interest. As such, TWAS identifies candidate genes for a trait, as well as the likely cell/tissue contexts. TWAS requires only summary statistics, making it easy to integrate eQTL and GWAS datasets from different cohorts. Because of these benefits, TWAS has become widely used to convert GWAS associations into candidate genes [6]. The TWAS framework is also applicable to other molecular traits, such as RNA splicing, or chromatin features, further broadening its utility [7].

A central question in TWAS analysis is whether the identified genes necessarily have causal effects on the phenotype. A simple analysis suggests this is not always the case (Fig. 1A). In one scenario, a non-causal gene *X*, has an eQTL *G*, that is in linkage disequilibrium (LD) with the eQTL of a nearby causal gene *X*′. This creates a non-causal association of the genetic component of *X* with the trait. In another scenario, *G* is in LD with a nearby causal variant, *G*′, which acts on the trait directly, again creating a non-causal association of the genetic component of *X* with the trait. In all these cases, false positive findings result from association of *G* with the trait, through pathways other than the gene *X*. This is known as “horizontal pleiotropy” in the literature, and is a key challenge that reduces the utility of TWAS [6].

**Figure 1.**
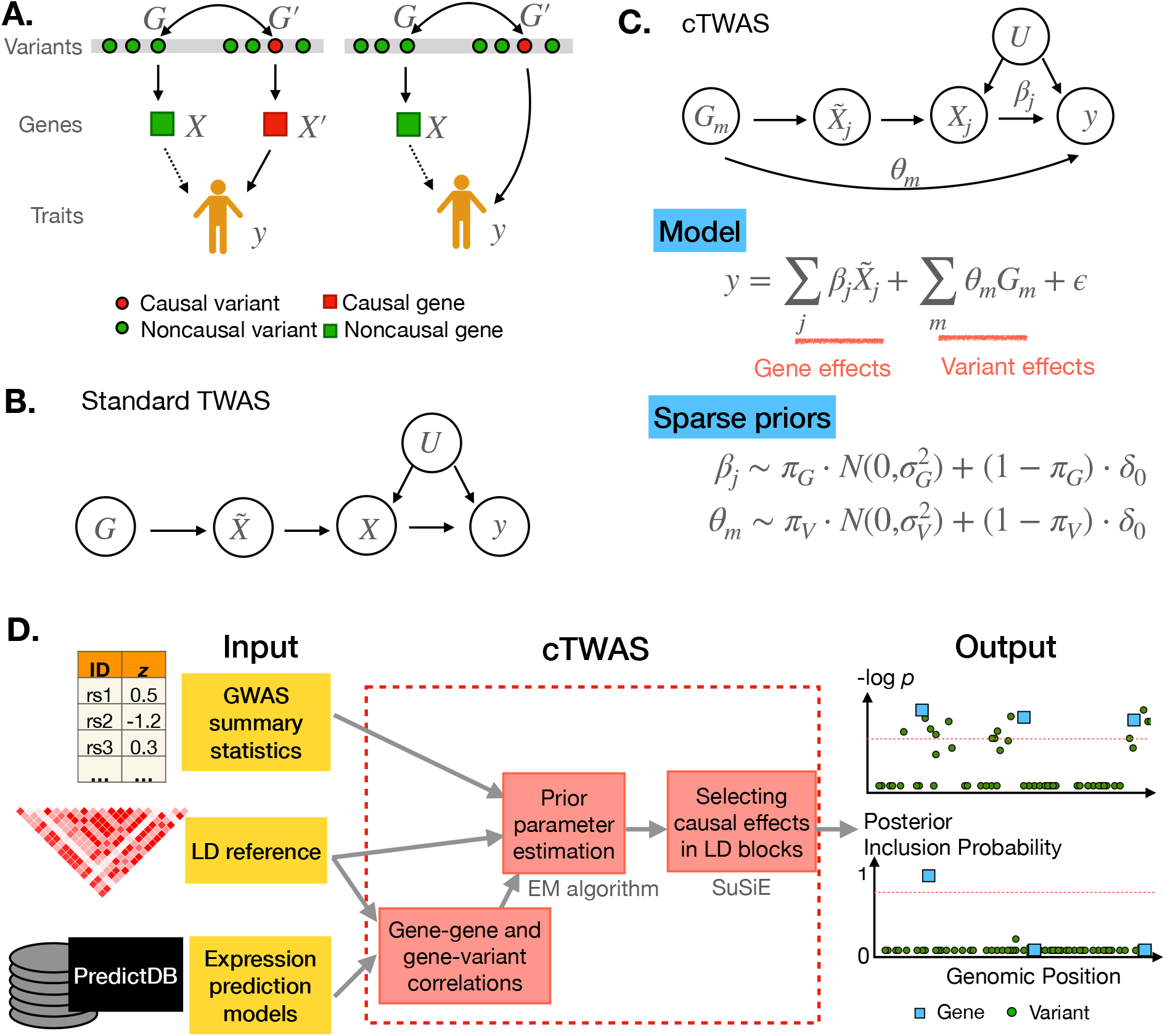
Overview of the cTWAS method. **A**. Two scenarios that violate the assumptions of TWAS and lead to false positive findings. *X* is a non-causal gene of the trait *y. X* is associated with the trait, shown as dashed arrows, because of the LD between its eQTL and nearby causal variants. Double-headed arrows represent LD between variants. **B**. The causal diagram implicitly assumed by TWAS. 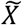 represents the cis-genetic component of gene expression *X. U* represents an environmental confounder. **C**. The model of cTWAS. *G*_*m*_ the genotype of the *m*th variant; 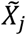 and *X*_*j*_ imputed expression and actual expression of the *j*th gene; *θ*_*m*_, direct effect of the *m*th variant on the trait *y*; *β*_*j*_, effect size of the *j*th gene; *ϵ*, error term; *δ*_0_, point mass at 0. *π*_*G*_ and *π*_*V*_, prior probability of being causal for genes and variants, respectively; 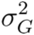 and 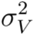, prior variance of the effect sizes of causal genes and variants, respectively. **D**. The workflow of the summary statistics version of cTWAS. The main steps of cTWAS are shown in the red boxes. cTWAS reports PIP for all genes and variants within LD blocks. *p* values for the genes and variants from marginal association tests are shown at the top as a comparison. Red dashed lines from the output panel indicate genome-wide significance level or the PIP threshold for genes.

Alternative methods have been proposed to jointly analyze eQTL and GWAS data, but they all face similar challenges. Colocalization methods test whether gene expression and a trait are affected by the same causal variant [8] [9]. Colocalization may alleviate the problems with TWAS, but does not fully resolve them. Indeed, in both scenarios described in Fig. 1A, if LD between the eQTL variant, *G*, and the nearby causal variant, *G*′, is strong enough, then *G* and *G*′ are effectively indistinguishable as potential causal variants of both gene expression and phenotype. False positives may also occur if the eQTL variant G has direct (pleiotropic) effects on the phenotype not through *X* [10]. Mendelian Randomization (MR) is another strategy to nominate causal genes, treating eQTLs of a gene as instrumental variables (IV) [11]. However, the potential pleiotropic effects of instruments and their LD with nearby causal variants, violate the key assumption of MR, leading to potential false positive findings. Several methods such as TWMR [12] and MR-JTI [13] attempted to address this issue by using a heterogeneity filter to remove variants that violate the MR assumption. But since most genes have only one or few cis-eQTLs (IVs), detecting heterogeneity is difficult in most cases. Lastly, methods such as FOCUS [14] and TWMR [12] jointly analyze multiple genes in a region. While these methods mitigated the challenge due to nearby genes (Fig. 1A, left), they largely failed to account for direct effects of nearby variants (Fig. 1A, right). Given that a large fraction of trait heritability is not mediated through eQTLs [15], this scenario is likely much more common and poses a bigger challenge.

Multiple lines of evidence suggest that the scenarios creating possible false positive findings are common. First, in TWAS, it is common to observe associations of multiple genes with a trait at a single locus, with most associations likely resulting from non-causal genes. Several examples were discussed in a perspective article on TWAS [6]. Colocalization analysis has a similar problem ; for example, in a Schizophrenia study, 282 genes in 78 risk loci, showed evidence of colocalization [16]. Secondly, at a biochemical level, co-regulation of genes by the same regulatory elements is common [17]. Thirdly, eQTLs are pervasive in the genome. In GTEx, half of all common variants are eQTLs in at least one tissue [18]. The implication is that chance associations (LD) between eQTLs of non-causal genes and causal variants are probably common [19]. Finally, while eQTLs are often associated with phenotypes, they mediate only 10-20% of heritability of complex traits [15]. This suggests that associations of eQTLs and a trait are often not due to causal effects of gene expression on the trait, but due to LD. All this evidence points to a critical need for better control of false discoveries in TWAS and other eQTL-based analysis.

Here we propose a new statistical framework to address the limitations of existing methods. Our approach can be viewed as a generalization of TWAS, which we term “causal-TWAS” (cTWAS). The fundamental problem of TWAS is that the imputed expression of any gene (the “focal gene”) is often correlated with expression of nearby genes or with other nearby variants (Fig. 1A). If these nearby genes and variants affect the trait, they induce a correlation between the trait and imputed expression of the focal gene. That is, they act as “confounders”, when testing association of the focal gene and the trait. We refer to these confounders as “genetic confounders” to distinguish them from the environmental confounders that are a common focus in the literature of MR.

This reasoning suggests a conceptually simple solution: when assessing association between the focal gene and trait through a regression model, we should include all genetic confounders in the model. In practice, implementing this strategy is complicated by high correlations that often exist among all these variables, which creates an identifiability challenge. Our key intuition is that causal signals in a genomic region, whether via gene expression or variants, are likely sparse. This motivates a Bayesian variable selection model, which has been widely used in statistical fine-mapping [20, 21, 22, 23]. Our approach, cTWAS, generalizes standard fine-mapping methods by including imputed gene expression and genetic variants in the same regression model. In simulations using realistic genetic parameters and applications to real data, cTWAS greatly reduces the number of false discoveries from TWAS, colocalization and MR-based methods, laying a foundation for reliable discovery of causal genes from GWAS.

## Results

### Overview of the cTWAS model

We start with a formal description of standard TWAS, using the language of causal diagrams [24]. We assume genetic variants, denoted as G, affect expression of a gene, *X*, which affects a trait *y* (Fig. 1B). Both *X* and *y* could be affected by unobserved environmental variable(s) *U*, such as diet. We introduce 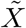, to denote the *cis*-genetic component of *X*. Importantly, the genetic variants *G* act on *y* only through 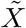. Under this model, the regression coefficient of 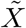, with *y* as the response, would give the causal effect of *X* on the trait. The confounder *U* is not a concern here because *U* contributes only to the non-genetic component of *X*, and thus does not create an association between 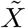 and *y*. In a formal language, the potentially non-causal path from 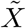 to *y* through *U*, has a collider, *X*, which blocks the association. Following similar analysis, we can see that TWAS results are also robust to “reverse causality” where *y* affects *X* (Fig. S1). In such a case, *X* is a collider in the paths from 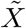 to *y*, making the two uncorrelated.

Unfortunately, the key assumption underlying TWAS, that *G* is not associated with *y* through other paths, is often violated. In the example discussed (Fig. 1A), 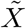 may become correlated with a nearby variant *G*′, or the genetic component of a nearby gene *X* ′. These confounders would create non-causal associations, technically known as backdoor paths, between 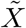 and *y*, leading to possible false discoveries by TWAS (Fig. S1).

To control for all potential confounders, cTWAS jointly models the dependence of phenotype on *all* imputed genes, and *all* variants, with their effect sizes denoted as *β*_*j*_ for genes and *θ*_*m*_ for variants, respectively (Fig. 1C). Joint estimation of all these parameters would then lead to causal effect estimates. In practice, to simplify computation, we partition the genome into disjoint blocks, with imputed expression and variants independent across blocks, and perform the analysis block-byblock. This is valid because in analysis of one region, the variants and imputed genes in other blocks are uncorrelated with the variables in the regression model, thus they would not create confounding.

The potentially high correlations among the variables in cTWAS poses a new challenge. Indeed, imputed expressions are calculated from eQTLs, thus they are often highly correlated with these eQTL variants. To address this challenge we assume that in any genomic region, causal effects, whether they are from genes or variants, are sparse. The problem then becomes similar to standard fine-mapping, where one aims to identify a small number of likely causal variants among many correlated ones. Additional intuitions help explain that the model can potentially learn gene effects despite collinearity. While most variants are non-functional, gene expression traits should be more likely to have causal effects *a priori*. Also, a causal gene may have multiple eQTLs, each of which would be associated with the trait. Thus, a single gene effect would be a more parsimonious explanation of data, compared with several independent variant effects.

We thus fit cTWAS using the statistical machinery developed for fine-mapping. We assume sparse prior distributions of the gene and variant effects (Fig. 1C), with an Empirical Bayes strategy to estimate these prior parameters from all data. Learning different prior parameters for genes vs. variants allows the model to favor gene effects over variant effects. With the estimated prior parameters, we infer likely causal genes and variants in each block, using SuSiE, a state-of-art method for fine-mapping [20] [21]. The final results of cTWAS are expressed as Posterior Inclusion Probabilities (PIPs) for genes and variants, representing the probabilities that genes or variants have non-zero effects (Fig. 1D). While cTWAS is formulated in terms of individual level data, we have also derived a version based on summary statistics (Fig. 1D, also see Methods).

The cTWAS model provided a general framework for joint analysis of eQTL and GWAS data. It generalizes and unifies a number of existing methods (see Discussion). We show that under a simple scenario where genes have only single eQTL variants, cTWAS reduces to existing colocalization methods (Supplementary Notes). Similar to TWAS, cTWAS can also be viewed as a two-stage MR method [25], where *cis*-genetic expression is used as the Instrumental Variable. But cTWAS accommodates horizontal pleiotropy through the inclusion of effects from variants and other genes. Lastly, while our primary goal is gene discovery, the learned prior parameters for gene and variant effects also allow us to estimate the proportion of heritability attributable to gene expression. This application of cTWAS is related to several other methods [15] [26].

### cTWAS controls false discoveries in simulation studies

We designed a realistic simulation setting to assess the performance of cTWAS. Simulations in previous studies often simulate individual regions, and these regions would usually contain causal genes. In our simulations, we created genome wide data across all regions, under realistic genetic parameters from previous stuides [15]. In particular, the percent of heritabilities mediated through eQTLs are low, so many regions may have causal variants, but not causal genes. Specifically, we used genotype data of common SNPs (minor allele frequency > 0.05) from ∼45k samples with white British ancestry from the UK biobank [27], and imputed gene expression using the prediction models from GTEx by FUSION [5]. We varied prior probabilities for genes and SNPs being causal, and prior effect size variances, with a total of 10 settings. We focused on two representative settings in the main results here, where the trait heritability is set at 0.5, and the percent of heritability explained by gene expression set at 10% (high gene PVE setting, where PVE stands for “percent of variance explained”), or 4% (low gene PVE). With these prior parameters, we sample causal genes, causal variants and their effect sizes, and then simulate traits according to our model (Fig. 1C) (Methods).

We first assessed the accuracy of parameter estimation. cTWAS estimated parameters were generally close to true values (gene results in Fig. 2A, and variant results in Fig. S2). In practice, what matters most is the ratio of prior probability of gene effects to that of variant effects. This “enrichment” parameter determines the extent to which the model favors gene effects vs. variant effects. Although cTWAS slightly under-estimated some prior parameters under some settings, the estimated enrichment remains accurate (Fig. 2A). Finally, cTWAS accurately estimated the proportion of trait variance explained by the gene effects (Fig. 2A).

**Figure 2.**
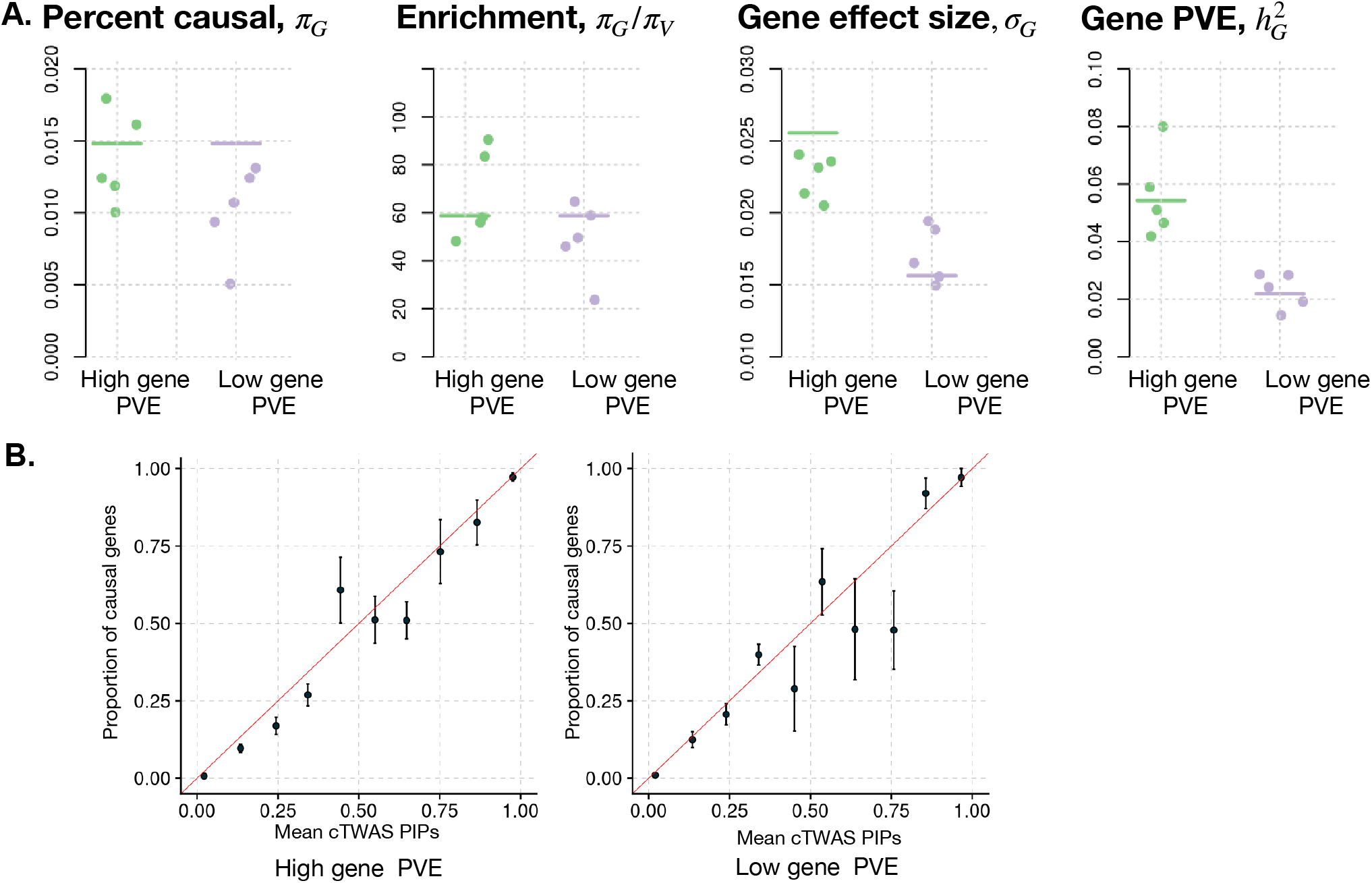
Parameter estimation and PIP calibration in simulations. **A**. Accuracy of the estimated parameters related to gene effects. Each plot shows one parameter: *π*_*G*_, prior probability for a gene being causal; enrichment (*π*_*G*_/*π*_*V*_), where *π*_*V*_ is prior probability for a variant being causal; effect size for gene; and PVE of gene. Results from two simulation settings are shown, the high and low gene PVE settings. Each dot represents the result from one out of five simulations. Horizontal bars show the true parameter values. **B**. Gene PIP calibration. Gene PIPs from all simulations are grouped into bins. The plot shows the proportion of true causal genes (Y axis) against the average PIPs (X axis) under each bin. A well calibrated method should produce points along the diagonal lines (red). +/- standard error is shown for each point in the vertical bars.

We also found that PIPs of genes computed by cTWAS are well-calibrated (Fig. 2B and Fig. S2). Good calibration means that at PIP > 0.9, we would expect at least 90% of genes above the threshold to be causal genes. Calibration is especially good at the high PIP range (90% or higher), which is what matters most in practice.

We found cTWAS successfully removed a large number of non-causal genes with highly significant associations in standard TWAS (Fig. 3A). We systematically compared the performance of cTWAS with other methods, including the standard TWAS implemented by FUSION [5], colocalization method (coloc) [28], MR-based methods (SMR with HEIDI filter [11], MR-JTI [13], and PMR-Egger [25]), and FOCUS [14], a multi-gene analysis method. Despite using stringent statistical thresholds, all these methods suffered from high false positive rates (Fig. 3B). Importantly, cTWAS controlled false positive rates with only a modest reduction of the number of discovered causal genes (Fig. 3B). Somewhat unexpectedly, the MR-based methods performed similarly or worse than other methods. We thus performed additional investigation of false positives in one of these methods, PMR-Egger (Supplementary Notes).

**Figure 3.**
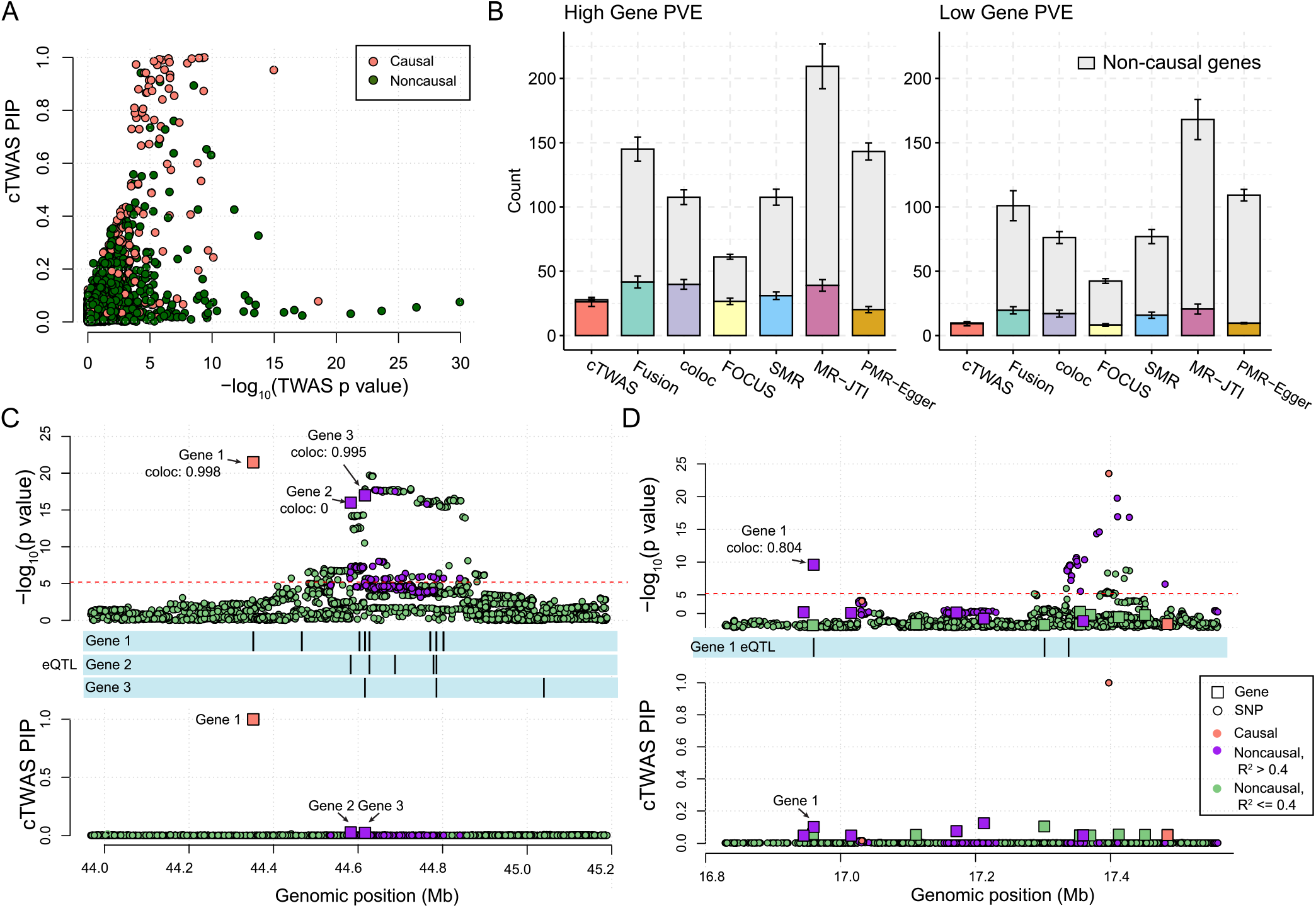
Comparison of cTWAS with other methods in simulations. **A**. Comparison of the p-values from standard TWAS and gene PIPs from cTWAS. The results were from one simulation run (parameters: gene PVE 0.052, gene prior 0.025, SNP PVE 0.50, SNP prior 0.00025). Each dot represents a gene and is colored based on whether it is a causal gene. **B**. Number of genes identified by various methods. We used the following significance thresholds for each method: cTWAS, PIP > 0.8; FUSION, Bonferroni corrected *p* < 0.05; coloc, PP4 > 0.8; FOCUS, PIP > 0.8; SMR, FDR < 0.05 and HEIDI *p* < 0.05; MR-JTI, Bonferroni corrected p < 0.05 for genes passing FDR < 0.05 from FUSION; and PMR-Egger, Bonferroni corrected *p* < 0.05. We use different colors for causal genes identified by each method, and the top gray bars indicate noncausal genes. **C** and **D**. Examples of how cTWAS avoided false positive genes. Top panel: -log_10_ p-values of genes (from TWAS) and SNPs in a region. Bottom panel: PIPs of genes and SNPs. Genes are represented by squares, with positions determined by transcription start sites, and SNPs represented by circles. Colors indicate whether the gene or SNP is causal (orange), noncausal but in LD with a causal effect (R^2^ between SNP genotype or imputed expression > 0.4, purple), or noncausal and not in LD with causal effect (green). The eQTLs of the genes are plotted in middle tracks. Transcriptome-wide significance cut off for TWAS was indicated by the red dotted line. In the top panel, the values of PP4 (probability of colocalization) from coloc analysis were shown for each gene of interest.

We illustrated, with two examples, how cTWAS was able to remove false positives. In the first example, the region has a single causal effect in Gene 1. However, because of LD between Gene 1’s eQTLs and eQTLs of other genes, two non-causal genes (Genes 2 and 3) also showed strong associations with the trait (Fig. 3C, top). cTWAS correctly identified Gene 1 as the true signal, and assigned low PIPs to the two other genes (Fig. 3C, bottom). Interestingly, coloc assigned high probability of colocalization to the non-causal Gene 3 (coloc PP4 = 0.995), probably due to LD between its eQTL with the eQTL of Gene 1. In the second example, the causal signal in the region is a SNP variant, but this SNP is in LD with the eQTL of Gene 1, creating a significant association of Gene 1 with the trait (Fig. 3D, top). By including both SNPs and genes in the model, cTWAS was able to correctly identify the SNP effect as the causal signal and assigned low PIP to Gene 1 (Fig. 3D, bottom). Coloc again gave high proprobality of colocalization to Gene 1 (PP4 = 0.8).

Finally, we investigated whether cTWAS is robust to model mis-specification. We sampled the effect sizes of causal genes and variants from mixtures of several normal distributions. The mixture distributions better capture the “long tails” of effect size distributions, i.e. some genes or SNPs have especially large effect sizes. We found that the gene effect enrichment was still accurately estimated, and PIPs were well calibrated (Fig. S3).

### cTWAS accurately identified causal genes of LDL cholesterol

We applied cTWAS to GWAS summary statistics of low density lipoprotein (LDL) cholesterol from the UK Biobank [29], which contains association z-scores of ≈8.7 million biallelic variants with MAF>0.01. We used the expression prediction models from GTEx [30] liver in PredictDB [4] [31]. After harmonizing eQTL data with the UK Biobank in-sample LD reference panel (see Methods), we included 9,881 protein-coding genes in analysis. cTWAS estimated that genes were over 62 times more likely than variants to be causal for LDL *a priori* (Fig. S4A). Genetic variants and imputed expression together explained 5.6% of variation of LDL (total heritability), 22.7% of which was attributable to expression. These estimates are in line with the 8.3% estimate for total heritability using LD score regression [32] and 33.5% of mediated heritability through expression using MESC [15]. The somewhat lower estimates of heritability from cTWAS likely resulted from the assumption of sparse causal effects.

Using the estimated prior parameters, cTWAS identified 35 genes with PIP > 0.8 (Table S1). In contrast, standard TWAS identified 215 genes at a Bonferonni-corrected significance threshold of 0.05. To assess these results, we used 69 previously-identified LDL-related genes as “silver standard” [13] [33] (see Methods). Following earlier work [34], we assessed the TWAS and cTWAS by their ability to distinguish silver standard genes from other genes within 1Mb, referred to as “bystanders”, most of which are probably false positives. We limited our analysis to 46 imputable genes out of 69 silver standard genes and 539 imputed bystander genes (Table S2). Among these genes, eight have PIP > 0.8 in cTWAS, and six of them are in the silver standard, leading to a precision of 75% (Fig. 4A). Standard TWAS, on the other hand, has a precision of 31% (19 out of 61).

**Figure 4.**
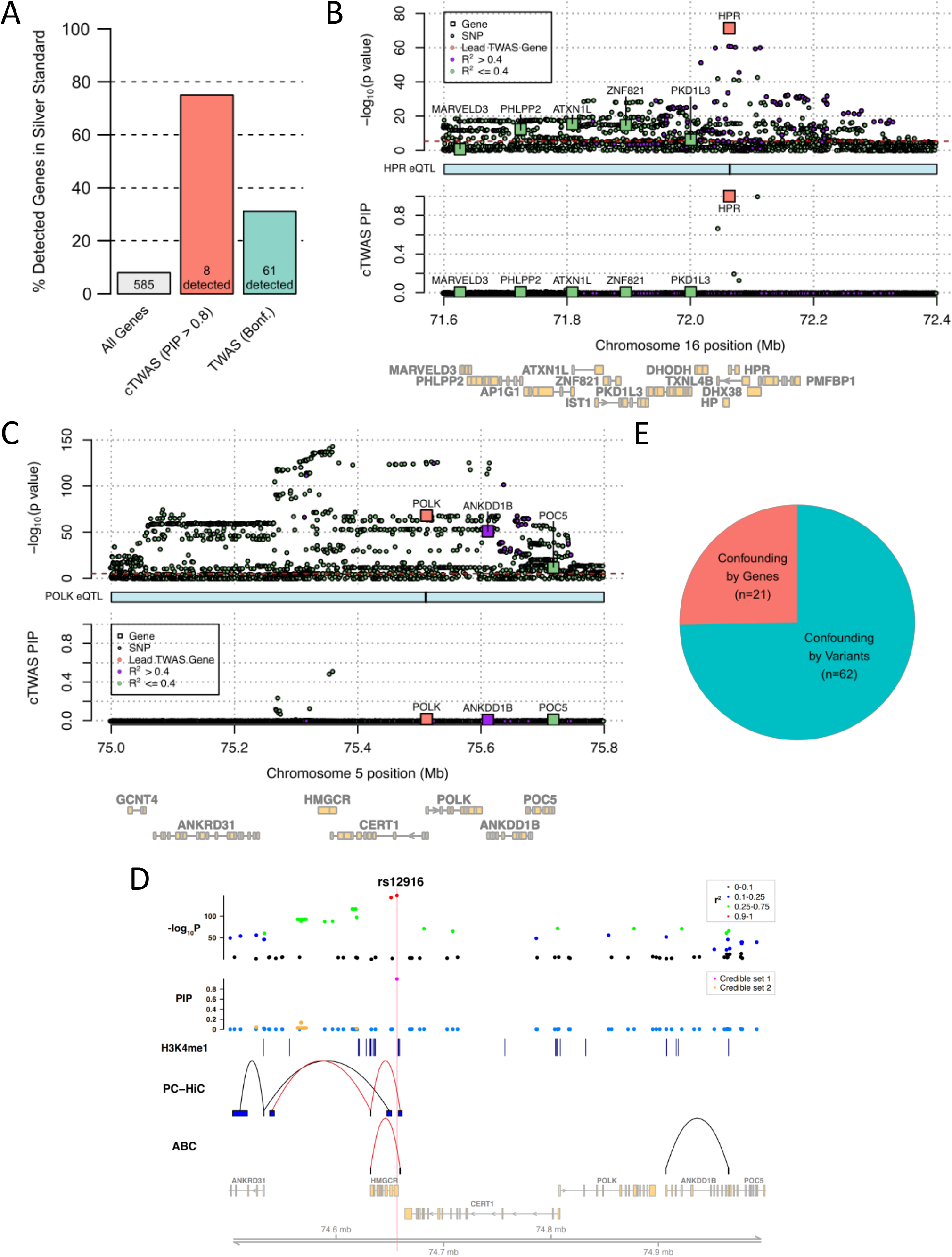
cTWAS avoids false discoveries of candidate genes of LDL cholesterol. **A**. Precision of cTWAS and TWAS in distinguishing LDL silver standard genes from nearby bystander genes. **B**. cTWAS results at the *HPR* locus. The top panel depicts the -log_10_ p-value of variants from GWAS and genes from TWAS. Each square represents a gene with position determined by its transcription start site. Each circle represents a variant. Colors indicate LD between the lead TWAS gene (orange) and nearby genes and variants: high LD (purple; R^2^ > 0.4) and low LD (green, R^2^ <= 0.4). The red dotted line indicates the transcriptome-wide significance threshold for TWAS (Bonferroni corrected *p* value < 0.05). The middle track represents the positions of the eQTL for the focal gene. The bottom panel represents the cTWAS PIPs for variants and genes at this locus. **C**. cTWAS results at the *POLK* locus. Figure legend is the same as in Figure 4B. **D**. Fine-mapping for the locus around *HMGCR* and *POLK* genes. The top two tracks represent the -log_10_ p-value of variants (with color representing LD with the lead variant) and their PIPs from fine-mapping with PolyFun-SuSiE (with color representing credible sets). Only variants with reported PIPs were shown in the plot. The third track represents liver H3K4me1 peak calls from ENCODE. The fourth track shows interactions identified from liver promoter-capture HiC (PC-HiC) data. The fifth track shows interactions identified from liver activity-by-contract (ABC) data. The links in red highlight regions looped to the *HMGCR* promoter. **E**. Sources of confounding for TWAS false positive findings. A TWAS gene was considered false positives if its cTWAS PIP ≤ 0.5. Only genes that can be assigned to a credible set were included in the analysis.

To illustrate how cTWAS avoided false positives, we examined two cases in detail. The first locus contains six genes significantly associated with LDL by TWAS, including *HPR* and five other genes (*ATXN1L, ZNF821, PHLPP2, PKD1L3*). cTWAS identified a single candidate gene, *HPR* (PIP = 1.000), while giving no evidence (PIP < 0.01) to all other genes (Fig. 4B). This can be explained by the fact that once controlling for *HPR*, cTWAS found no additional signal in other genes. Literature evidence suggests that *HPR*, haptoglobin-related protein that binds hemoglobin and apolipoprotein-L [35], is the likely causal gene at this locus. This example illustrates that cTWAS avoids false positives due to confounding with nearby gene expression.

The second locus has three genes strongly associated with LDL by TWAS: *POLK, ANKDD1B*, and *POC5* (Fig. 4C, top). A recent method, MR-JTI [13], has highlighted *POLK*, DNA polymerase κ, as the potential causal gene at this locus, and proposed a connection between DNA repair and regulation of LDL. The associations of the three genes, however, were much weaker than some nearby variants (Fig. 4C, bottom). This pattern was reflected in the results of cTWAS. cTWAS selected several highly associated variants as causal signals, while giving little evidence to all three genes. This suggests that the associations of the genes with LDL are due to their correlations with true causal variants, which may act through other pathways.

To better understand these results, we inspected variant-level fine-mapping results from PolyFun, which uses functional information of variants to improve the accuracy of fine-mapping [36]. PolyFun identified two causal signals in the region, with the variants in the credible set all inside or close to the gene *HMGCR*, whose expression was not imputable in our data (Fig. 4D). All the credible set variants are far from the three TWAS genes, including POLK (>200 Kb). Additionally, promoter-capture Hi-C (PC-HiC) data and the Activity-by-contact (ABC) score in liver provided no evidence linking these variants to *POLK*. Instead, the top variant in the locus, rs12916 (PIP = 0.99) is within the 3’ UTR of *HMGCR*, and 1310 bp away from a chromatin loop interacting with the *HMGCR* promoter, in both PCHiC and ABC (Fig. 4D). Consistent with these results, *HMGCR* is the rate-limiting enzyme for cholesterol synthesis and the target of statin, a key drug for reducing LDL levels [37]. All the evidence thus points to *HMGCR*, instead of *POLK* or other genes, as the causal gene in this region. This example demonstrates that by controlling nearby genetic variants, cTWAS is able to avoid false positive genes.

We systematically evaluated the sources of possible false positive findings from standard TWAS. We call a gene a likely false positive if it is significant under TWAS (Bonferroni threshold), but PIP < 0.5 under cTWAS. These cases were classified as “confounding by genes” or “confounding by variants”, depending on whether the low PIPs of these genes were primarily driven by nearby genes or variants (see Methods). The majority of the 83 false positive genes analyzed (74.7%, Fig. 4E) were driven by confounding variants. These results show that the greatest risk of TWAS is not co-regulation of genes, i.e. shared eQTLs among nearby genes, but rather the correlation of genes with nearby variants whose effects are not manifested as eQTLs.

To seek new insights into the genetics of LDL, we evaluated the functions of 35 genes with cTWAS PIP > 0.8 (Fig. 5A). Only six of these genes were in the curated silver standard genes, and 20 of 35 genes were not the nearest genes of GWAS lead variants (Fig. 5A). The 35 genes were enriched for multiple cholesterol-related Gene Ontology (GO) Biological Process terms (FDR < 0.05) (Fig. 5B, 13 non-redundant terms shown; Table S3). To assess the novelty of these GO terms, we performed GO enrichment analysis of silver standard genes (Table S4) and GWAS gene set analysis using MAGMA (Table S5). Several GO terms enriched in cTWAS genes were not found in GO terms from the silver standard or MAGMA analysis, including “peptidyl-serine phosphorylation” and “activin receptor signaling pathway”, highlighting the importance of signal transduction in LDL regulation. Activin signaling, in particular, regulates metabolic processes including lipolysis and energy homeostasis [38] [39]. The cTWAS genes associated with the two terms include well-known LDL genes, such as *CSNK1G3, TNKS*, and *GAS6*, as well as novel and promising genes such as *ACVR1C*, an activin receptor, and *PRKD2* (Fig. 5C; Fig. S4B; see Discussion). In the cases of both *ACVR1C* and *PRKD2*, no nearby variant reaches genome-wide significance, thus the genes would be missed by standard GWAS analysis.

**Figure 5.**
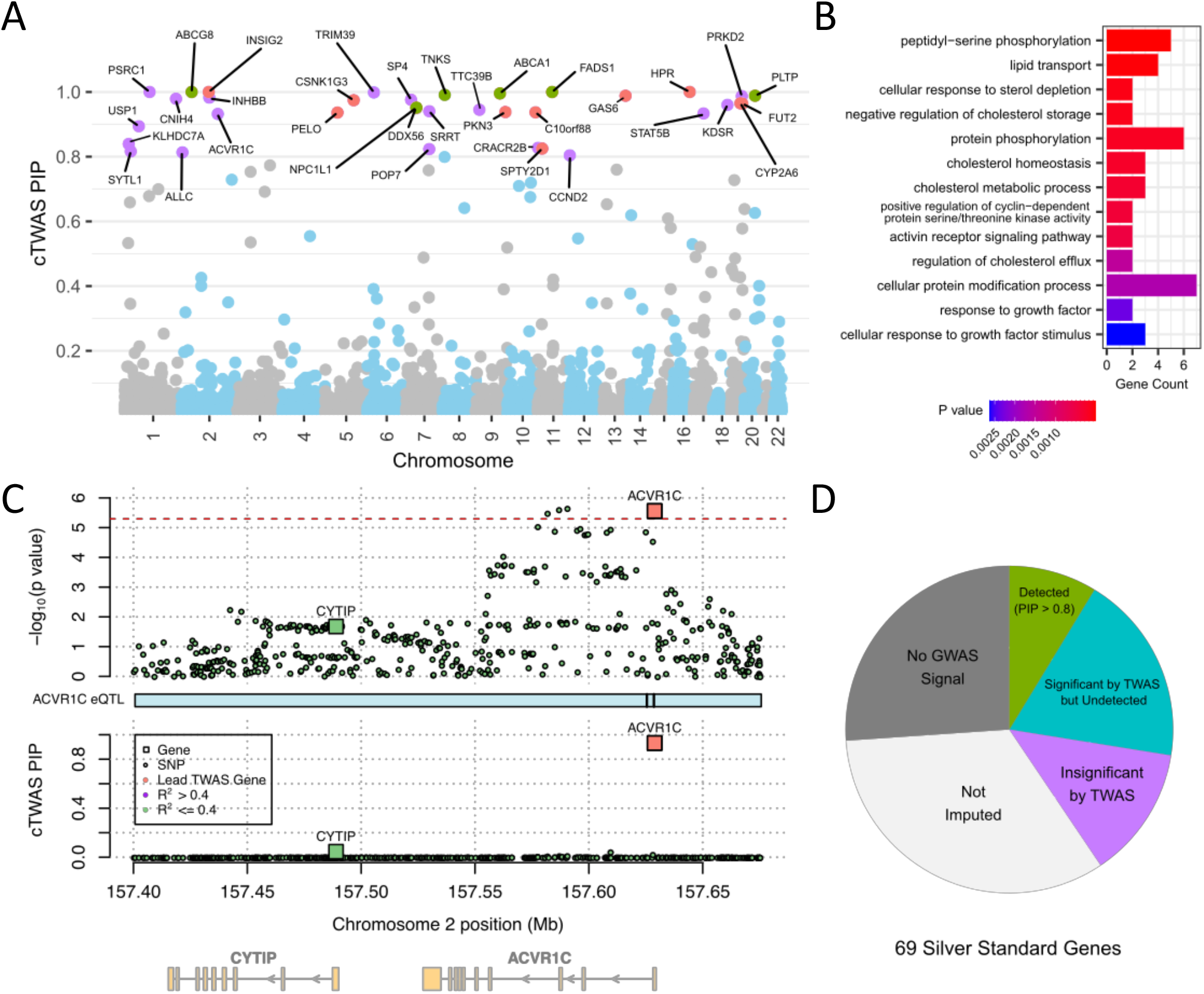
Candidate genes and pathways for LDL discovered by cTWAS. **A**. PIPs of 9,881 liver genes from cTWAS analysis of LDL cholesterol. The labeled genes have PIP > 0.8, colored based on existing evidence for LDL function: silver standard genes (green); nearest genes of genome-wide significant loci but not in silver standard (orange); or otherwise novel (purple). **B**. GO Biological Process terms enriched (FDR < 0.05) among 35 detected LDL genes, at PIP > 0.8. Redundant terms were omitted for clarity (see Methods). Gene count means the number of detected genes associated with a GO term. **C**. cTWAS results at the *ACVR1C* locus. Figure legend is the same as in Figure 4B and 4C. **D**. Summary of cTWAS outcomes for all 69 silver standard genes, into the following categories: detected by cTWAS at PIP > 0.8 (“Detected (PIP > 0.8)”); significant by TWAS (Bonferroni threshold), but not detected by cTWAS (“Significant by TWAS but Undetected”); insignificant by TWAS and not detected by cTWAS, with a genome-wide significant GWAS variant within 500kb (“Insignificant by TWAS”); genes have no expression prediction models (“Not imputed”); not detected by cTWAS and TWAS, and no genome-wide significant GWAS variant within 500kb (“No GWAS Signal”).

While cTWAS reduced false positives and identified promising LDL candidate genes, its power seemed low. Out of 69 silver standard genes, 6 (8.7%) have PIP > 0.8 under cTWAS. To understand why, we categorized the outcome of cTWAS for all 69 genes (Fig. 5D). Many silver standard genes had no significant GWAS association signals nearby (26.1%, 18 of 69), no imputable liver expression (33.3%, 23 of 69), or insignificant TWAS associations (13.0%, 9 of 69). These results suggest that to improve the power of cTWAS, and eQTL-based methods in general, it is necessary to improve the power of GWAS, the power of eQTL studies, and include more trait-related tissues/cell types (see Discussion).

### cTWAS discovered candidate genes of several common traits

We applied cTWAS to several commonly studied traits, including Inflammatory Bowel Disease (IBD), Systolic Blood pressure (SBP) and Schizophrenia (SCZ). We used the expression prediction models of protein-coding genes from PredictDB [40, 31, 4] across 49 tissues in GTEx [30]. These models borrowed information across tissues to improve prediction accuracy in any specific tissue [41]. The numbers of imputed protein-coding genes across tissues ranged from 6,591 to 11,985 (Fig. S5). We ran cTWAS analysis for a trait in each tissue separately. We summarized the results below, with a particular emphasis on IBD as a representative trait.

We first assessed the parameters learned by cTWAS. The prior probability of an imputable gene being causal ranged from 0.17% to 2.16% across tissue-trait pairs (Fig. S6, Table S6). For example, for IBD, the top tissue is whole blood, with the percent of causal genes 1.54%. The estimated proportions of heritability of traits explained by the genetic components of expression were generally small, e.g. for IBD, they range from 4% to 15% (Fig. 6A, Table S6). These estimates were generally in line with estimated values from MESC (Fig. 6A, Fig. S7, Table S7). Altogether, these results highlight the ability of cTWAS to infer important genetic parameters.

**Figure 6.**
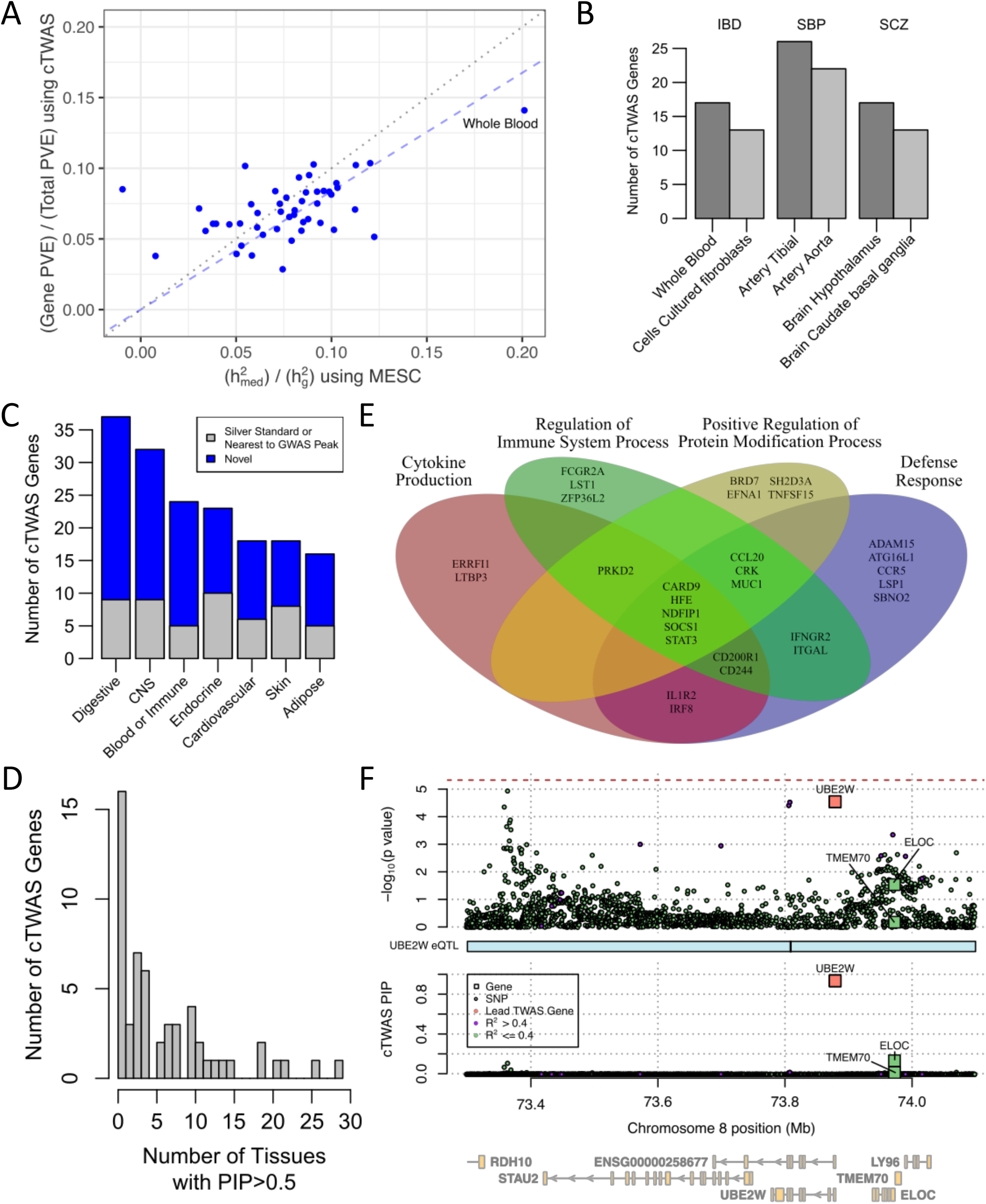
cTWAS analysis of IBD and other traits using all GTEx tissues. **A**. Estimated percentages of heritability mediated by eQTLs for IBD using MESC (X-axis) and cTWAS (Y-axis). Each dot represents the result from one tissue. The black dotted line denotes equivalence between the methods, and the blue dashed line denotes the slope relating cTWAS and MESC mediated heritability estimates. **B**. The number of cTWAS genes detected at PIP > 0.8 in the top two tissues for IBD, SBP, and SCZ. Top tissues per trait are determined by the number of detected genes. **C**. The number of cTWAS genes detected for IBD at PIP > 0.8 across major tissue groups. A gene was detected in a tissue group if it was detected in any of the tissues in that group. A gene was considered novel if it was not a silver standard gene (“Known”) or if it was not the nearest gene to a genome-wide significant locus for IBD (“Nearest”). **D**. The number of tissues with cTWAS PIP > 0.5 for 56 IBD genes detected at PIP > 0.8 in the “Blood/Immune’’ or “Digestive” tissue groups. **E**. Non-redundant GO terms enriched among 56 detected IBD genes in the “Blood/Immune” or “Digestive” tissue groups. These terms were found using the Weight Set Cover method from WebGestalt. **F**. cTWAS results for IBD at the *UBE2W* locus using the “Colon Transverse” eQTL data. Figure legend is the same as the previous locus plots.

We next assessed the number of high confidence genes, at PIP > 0.8, found by cTWAS (Fig. S8). Among the top 2 tissues per trait, cTWAS identified 13 - 26 genes (Fig. 6B). In general, the number of cTWAS genes were much smaller than those from standard TWAS (Fig. S9). For instance, for IBD, while TWAS reported 68-125 genes across 49 tissues, cTWAS identified 0-17 genes (Fig. S9). These results confirm that only a small proportion of genes found by TWAS are likely causal genes.

Given that relatively few genes were discovered in each tissue separately, we grouped related tissues into “tissue groups” and took the union of genes across all tissues within a group (Fig. 6C, Fig. S10). The top tissue groups include trait-relevant tissues, such as “Digestive” tissue for IBD (Fig. 6C), “Cardiovascular” for SBP and “Central Nervous Systems” for SCZ (Fig. S10). The number of discovered genes in the top tissue group per trait ranged from 37 (IBD) to 48 (SBP), highlighting the increased power of discovery from multiple tissues. We also assessed the novelty of the found genes. In the case of IBD, most of cTWAS genes were not in the curated genes for IBD [42], and not the nearest protein-coding genes of lead genome-wide significant GWAS variants (Fig. 6C).

A natural question is whether the discovered genes are limited to specific tissues. We found that most cTWAS genes were identified in a small number of tissues (Fig. S11). For instance, for the 56 IBD genes found in the “Blood/Immune”, or “Digestive” tissue groups, 57.1% of them were found, at a relaxed threshold of PIP > 0.5, in 5 or fewer tissues(Fig. 6D). One caveat in interpreting these findings is that the power of discovery is low, so cTWAS may underestimate the number of tissues for discovered genes.

We examined specific genes found by cTWAS (Table S8 for all traits). We focused our analysis here on 56 genes found for IBD, at PIP > 0.8, in the two biologically relevant tissue groups: digestive and blood/immune (Table S9). At a more stringent PIP > 0.9, 30 genes were found (Table 1). The set of 56 genes included well-known IBD genes, such as *TNFSF15, CARD9, RNF186, ITGAL*, and *ATG16L1*. Gene set enrichment analysis revealed many IBD-related GO terms in the 56 genes (Table S10). Using Weighted Set Cover [43], a method to extract non-redundant GO terms, we identified four GO terms, such as “cytokine production”, and “defense response”. 29 out of 56 genes were associated with at least one of these four terms (Fig. 6E).

**Table 1.**
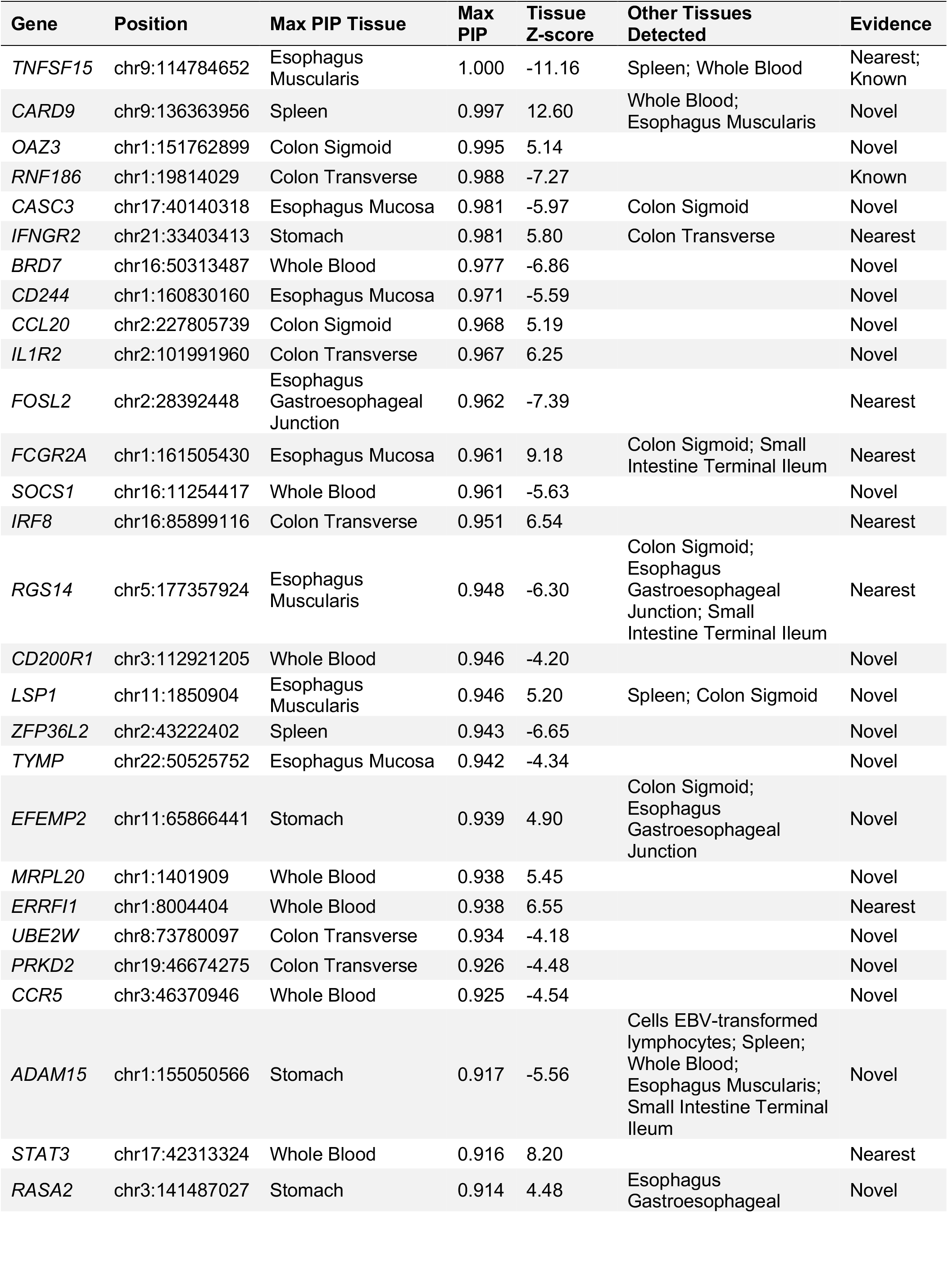
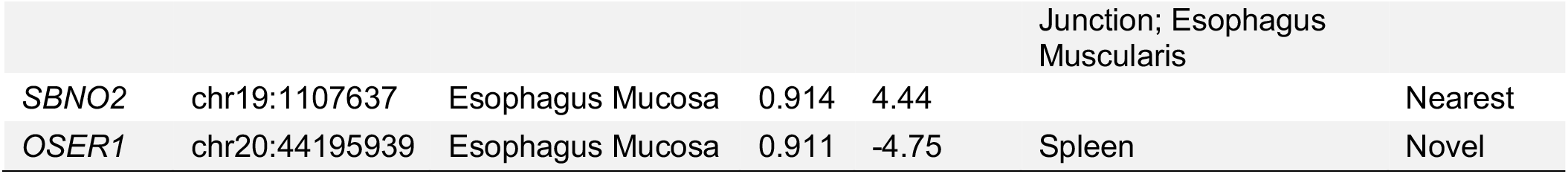
IBD genes detected by cTWAS in the Blood/Immune and Digestive tissue groups. Genes detected by cTWAS at PIP > 0.9 in at least one of the tissues in the two tissue groups. The Max PIP Tissue, Max PIP, and Z-score columns denote the tissue with the highest PIP for each gene, its corresponding PIP, and Z-score from TWAS in that tissue. The Other Tissues column lists any additional tissues with cTWAS PIP > 0.8. The Evidence column denotes whether each gene is in the silver standard gene list (“Known”); the nearest gene to genome-wide significant GWAS peak for IBD (“Nearest”); or otherwise novel (“Novel”).

We highlight some novel genes found by cTWAS. Many of these genes are located within known IBD-associated loci and have immune functions: *IFNGR2, FOSL2, STAT3, FCGR2A, IRF8*, and *ZFP36L2* (see Discussion). cTWAS also identified novel genes in the loci whose associations fall below standard GWAS cutoff. Some of these genes have IBD-related functions, including *UBE2W* (Fig. 6F), *TYMP, LSP1*, and *CCR5* (Fig. S12, see Discussion). For example, *UBE2W* is a ubiquitin-conjugating enzyme. Ubiquitination is a post-translational modification that controls multiple steps in autophagy, a key process implicated in IBD. Indeed, *UBE2W* knockdown mice showed mucosal injuries, and its overexpression ameliorated the severity of experimental colitis, a model of IBD [44].

## Discussion

Expression QTL data are commonly used to annotate the effects of genetic variants and to nominate candidate genes for complex traits. Existing methods for such analysis, however, are susceptible to finding non-causal genes. This happens because these methods fail to account for the pleiotropic effects of eQTLs or their LD with nearby causal variants. To address this challenge, we generalize the TWAS model by explicitly and jointly modeling the effects of all gene expression traits and genetic variants in a region. Through extensive simulations, we show that cTWAS achieves calibrated false discovery rates. In the analysis of several common GWAS traits, cTWAS discovered a number of candidate genes for these traits, highlighting its potential as a powerful gene discovery tool.

cTWAS is related to existing methods for integrating eQTL and GWAS data but has several key advantages. When the gene of interest has a single causal eQTL, and the gene is the only causal gene in a locus, cTWAS reduces to the colocalization models (Supplementary Notes) [28, 8, 45]. Despite this connection, cTWAS has key advantages over colocalization. Colocalization typically focuses on individual variants, yet cTWAS uses imputed gene expression, which combines the effects of multiple variants. While colocalization can be generalized to allow for multiple causal variants [46], it does not explicitly account for the combined effects of variants. cTWAS can also be viewed as a generalization of FOCUS, which uses a fine-mapping framework, similar to ours, but includes mostly gene effects, with a very simple model of variant effects. As others have pointed out [15], and our results showed (Fig. 4E, Fig. 6A), gene expression traits mediate a relatively small percent of trait heritability, so confounding by nearby variants is a much more common source of false discoveries.

cTWAS is also related to some MR-based methods. Similar to cTWAS, PMR-Egger [25] jointly models the effect of a gene on a phenotype, and potential pleiotropic effects of variants, following the Egger extension of MR. This model, however, analyzes one gene at a time, and its treatment of pleiotropy is overly simplified, assuming all genetic instruments of a gene have identical pleiotropic effects. TWMR [12] uses multivariate MR to jointly infer the causal effects of multiple genes in a locus. But it does not explicitly model the pleiotropic effects from variants. Instead, it relies on a heterogeneity filter to remove invalid instruments. Given that most genes have few independent cis-eQTLs/instruments, detecting heterogeneity would be difficult.

One main finding from our study is that TWAS false positive findings were often due to correlation of imputed expression with nearby causal variants (“confounding by variants”, Fig. 4E). These results reflect the fact that *cis*-genetic components of expression explain relatively low proportions of trait heritability [15] (Fig. 6A). Indeed, a recent study has made an evolutionary argument for this observation [47]. Namely, genetic variants with large effects on the risk genes of diseases are under strong selection and as a result, their frequencies would be low and their eQTL effects hard to detect. Despite these observations, we view eQTLs, combined with our discovery framework, as an important tool for disease gene discovery. The cTWAS genes were often found using prediction models that combine the effects of multiple, potentially weak, eQTLs, suggesting that cTWAS is less dependent on variants with large eQTL effects. Additionally, combining data across multiple tissues can increase power, as shown in the case of IBD (Fig. 6C). Given the strong community interest in profiling eQTLs under various conditions, and across multiple cell types [48], leveraging these data will likely substantially increase power to discover disease genes. Lastly, our method can be applied to other types of molecular QTL data, e.g. chromatin accessibility QTLs, which may explain a large fraction of heritability missed by eQTLs [7].

The application of cTWAS to several complex traits suggested interesting and novel candidate genes. In the case of LDL, we identified two genes, ACVR1C and INHBB, in the signaling pathway of activin, a signaling molecule in the TGF-beta superfamily. ACVR1C is a receptor of activin A and INHBB is a subunit of activin B. Both activin A and B regulate lipolysis in hepatocytes and adipocytes and alter lipid compositions [49] [38]. In an exome-wide association study, a loss-of-function variant of ACVR1C was associated with metabolic phenotypes such as triglycerides and high-density lipoprotein level [50]. We identified another signal transduction gene, a protein kinase, PRKD2, as a candidate gene affecting LDL. PRKD2 deficiency in mice triggers hyperinsulinemia, metabolic disorders and dysregulation of LDL [51].

Our study of IBD revealed a number of interesting candidate IBD risk genes (Table 1). IFNGR2 is a receptor of interferon-gamma, a proinflammatory cytokine, whose activation may result in intestinal lesions [52]. IRF8 is an important regulator of multiple immune processes ranging from antigen presentation to response to cytokines [53]. IRF8 deficiency in an experimental model of colitis resulted in more severe inflammation [54]. CCR5 is a chemokine receptor, best known for its role in HIV infection. CCR5 blockade in mice inhibited leukocyte trafficking and reduced mucosal inflammation in murine colitis [55]. TYMP encodes thymidine phosphorylase, and TYMP mutations cause Mitochondrial Neurogastrointestinal Encephalomyopathy (MNGIE), a rare recessive disease [56]. The disease causes gastrointestinal symptoms and is often misdiagnosed as celiac or Crohn’s disease [56]. LSP1 is a regulator of T-cell migration, and deletion of LSP1 gene has been implicated in rheumatoid arthritis, an autoimmune disease [57].

We discuss possible directions of further development of cTWAS. Firstly, it is relatively straight-forward to include more tissues or cell types in the cTWAS model. This can be done by including multiple groups of explanatory variables (imputed expressions), with different priors for different groups. Having a multi-tissue model may increase the power to detect causal genes and help identify the “causal tissue” for these genes. Secondly, we treated imputed expression levels as given. It may be helpful to account for imputation errors in the model [58]. Lastly, it would be interesting to generalize the model to allow joint analysis of multiple types of molecular QTL data.

In conclusion, by modeling genetic variants and imputed gene expression jointly, cTWAS accounts for pleiotropic effects and LD, creating a robust framework for detecting causal genes. With the large amount of molecular QTL datasets available and being generated, cTWAS promises to translate genetic associations of diseases into knowledge of risk genes, disease mechanisms and potential therapeutic targets.

## Code availability

Our software is available at https://xinhe-lab.github.io/ctwas/.

## Methods

### Model of individual level data

Let *y* be the quantitative phenotype, assumed to be standardized, of an individual. We assume that *y* depends on imputed gene expressions and variant genotypes of the individual. We denote *X*_*j*_ the expression of the gene *j*, 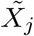 as its *cis*-genetic component, and *G*_*m*_ the genotype of the variant *m*. We assume that 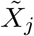 is given, imputed from a pre-trained expression prediction model, and the imputation errors/uncertainty would be ignored. We have the following regression model:

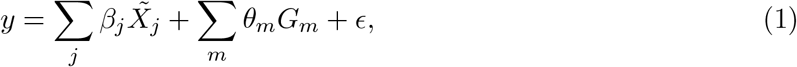

where *β*_*j*_ and *θ*_*m*_ are the effect sizes of gene expression *j* and the variant *m*, respectively. *ϵ* is an normally distributed error terms, i.e. *ϵ ∼ N*(0, *σ*^*2*^), and is assumed to be independent across individuals. In practice, we standardize both 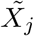 and *G*_*m*_ to make the variance equal to 1 for all the genes and variants.

To obtain the imputed expressions, we use existing expression prediction models. Specifically, the imputed expression of a gene *j* is defined as*∑*_*l*_ *w*_*jl*_*G*_*l*_, where *G*_*l*_ is the genotype of variant *l*, and *w*_*jl*_ is the weight of the *l*-th variant in gene *j*’s expression prediction model. We assume that these weights are given at the standardized scale, i.e. the weights were derived using standardized variant genotypes. This is the case for the FUSION expression models (http://gusevlab.org/projects/fusion/). When the provided weights are not on the standardized scale, for example, from PredictDB (https://predictdb.org/), these weights must be scaled. This can be done via multiplying the weights by genotype variances from the LD reference.

We specify different prior distributions of gene effects *β*_*j*_’s, and variant effects *θ*_*m*_’s. To describe these priors, we note that our model is a special case of a more general regression model, where explanatory variables come from multiple groups with different distributions of effect sizes. We write the general model with *K* groups of explanatory variables as:

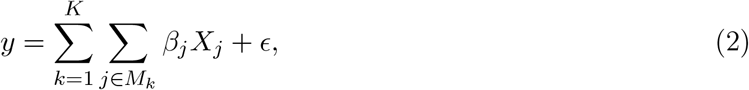

where *X*_*j*_ is *j*th explanatory variable and *j* ∈ 2 *M*_*k*_ denotes that it belongs to group *k*. In our case, the model has two groups of variables, imputed gene expressions and genetic variants. For simplicity of notations, we will use this general model in our following discussions. We assign a spike-and-slab prior distribution for the effect of variable *j*, with group-specific prior parameters. Specifically, when *j ϵ M*_*k*_, we denote *γ*_*j*_ an indicator of whether *X*_*j*_ has non-zero effect:

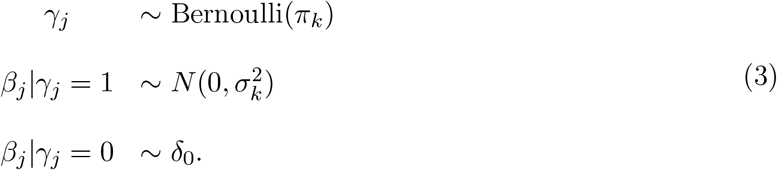

Here *δ*_0_ is the Dirac’s delta function, *π*_*k*_ = *P*(*γ*_*j*_ = 1|*j ϵM*_*k*_) is the prior probability of the *j*th variable from group *k* being casual to the trait (non-zero effect), and 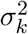 is the prior variance of the effect size of causal variables in the group *k*.

### Inference of the individual level model

The inference has two main steps. In the first step, we estimate the prior parameters 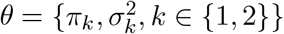 for the two groups, gene effects and variants effects. In the second step, we use the estimated *θ*, and compute the posterior inclusion probability (PIP) of each variable, defined as the posterior probability of *γ*_*j*_ =1 given all the data and parameters.

The parameter estimation is done by Maximum Likelihood. Let ***y***_*n*×1_ be the data of the response variable, where *n* is the sample size. Let ***X***_*n*×*p*_ = [***X***_1_ ***X***_2_ … ***X***_*p*_] be the data of all the *p* explanatory variables. The likelihood of our model is given by

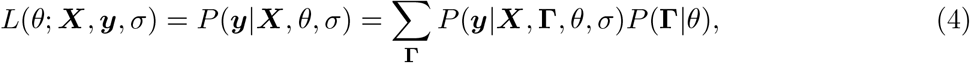

where **r** = [*γ, γ*, …, *γ*_*p*_] represents the “configuration” of the causal (non-zero effect) status of all variables. We note that *σ* is the standard deviation of the phenotypic variance, and is assumed to be given (see below). To maximize the likelihood, we use the Expectation-Maximization(EM) algorithm. In the E-step, we obtain the expectation of log likelihood over **r**, *𝔼*_**r**_log*P*(***X, y*, r**|*θ*^(*t*)^, *σ*), where *θ*^(*t*)^ is the parameter values in the *t*-th iteration. In the M-step, we update *θ*^(*t*)^ using the following rules to maximize the expectation from the E-step (see Supplementary Notes):

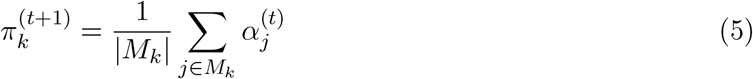

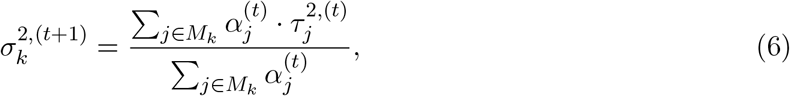

where |*M*_*k*_| is the number of variables in group *k*, 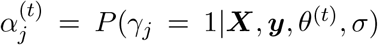 is the PIP of variable *j* given data and current parameter values *θ*^(*t*)^, 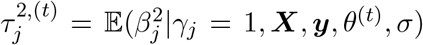 is the second moment of the posterior effect size of variable *j*, given that it is a causal variable. The update rules have simple interpretations. The new parameter 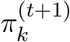 is simply the average PIP of all variables in the group *k* and the new 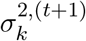 is the weighted average of the second moment of the posterior effect sizes.

Computing *σ*_*j*_ and 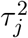 at the *t*-th iteration (we removed superscript *t* from now on for simplicity) using all variables in the genome is computationally challenging. To reduce the computational burden, we divide the genome into linkage disequilibrium (LD) blocks using LDetect [59] with variants approximately independent between blocks. We assign a gene into an LD block if all SNPs in its expression prediction model fall into that block. If the variants of the prediction model of any gene span multiple LD blocks, we merge all such blocks into a new block. We will then compute *σ*_*j*_ and 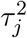 of the variables in each block independently, while still using all variables in the genome to update the parameters using Equations 5 and 6.

Even within a single block, there may still be hundreds to thousands of variables. This poses a challenge to compute *σ*_*j*_ and 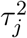, as it requires marginalization of **r**. To address this challenge, we first notice that our problem is now reduced to standard fine-mapping or Bayesian variable selection problem, with potentially different prior distributions of the effects of different variables. So we borrow from standard fine-mapping literature to compute *σ*_*j*_ and 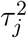 [20, 21]. We further assume that there is at most one causal variable per block (single effect assumption). So the only possible configurations are: **r**_*j*_, the configuration where *γ*_*j*_ = 1 and *γ*_*i*_ = 0 if *I* ≠ *j*, and **r**_0_, the configuration where none of the variables in this block is causal. We also assume that a single block explains a small amount of the variance of the trait, so the residual variance of the trait, *σ*^2^ would be approximately equal to 1, the total phenotypic variance (*y* is assumed to be standardized). With these assumptions, we have:

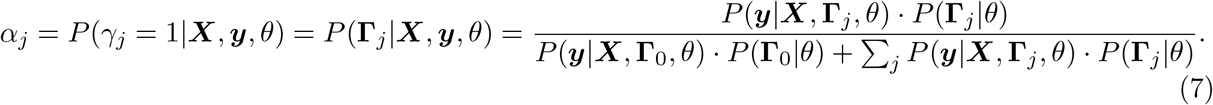

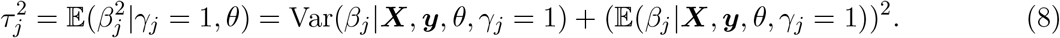

Both *α*_*j*_ and 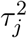 have closed forms under the single effect assumption. Specifically, the PIP of variable *j* (*α*_*j*_) is a simple function of prior inclusion probability *p*_*j*_ = *P*(*γ*_*j*_ = 1|*θ*), and the Bayes factors (BFs) of the variables:

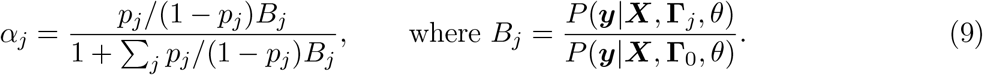

The BFs can be computed using the Wakefield formula [60]. See Supplementary Notes for details.

We provided some justifications of the single effect assumption. In statistical fine-mapping, several methods have been developed to incorporate functional information of variants to set the prior inclusion probabilities. In such models, it has been found that for the purpose of estimating parameters of the prior distributions, the single effect approximation is often sufficient [9]. Furthermore, in our implementation, we prune large blocks that are likely to contain multiple causal variables in the parameter estimation step (see Supplementary Notes). Finally, our simulations showed that the parameters estimated under single-effect assumption were generally accurate (Figure 2 of the main text).

After we estimate the prior parameters, we apply SuSiE [20], a fine-mapping method, on all variables, including both genes and variants, in each block. Note that all blocks, including the large blocks pruned in the parameter estimation step, will be analyzed. In applying SuSiE, we set the prior probability and prior effect variance of each variable, using the estimated parameters of the group (genes or variants) that this variable belongs to. We allow multiple causal variables by setting *L* =5 in SuSiE and assign null weight as 1 − *∑*_*j*_ *p*_*j*_. SuSiE will then return PIPs of all genes and variants in each LD block.

### Model of summary statistics

It is often easier to work with summary statistics from GWAS data. The summary data would include the effect size estimates of variants, 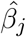, and their standard errors *s*_*j*_, as well as the LD between all pairs of variants, denoted as the matrix ***R***. The effect sizes can be standardized, denoted as 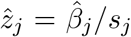. Given that the summary data has only variant information, our model would first need to expand the summary data to include gene information. Specifically, we compute the marginal association of each imputed gene with the GWAS trait, and the correlation of any gene with all other genes and all the variants. These calculations will be described below. Once computed, we will have marginal associations of all variables, including genes and variants, analysis. 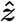, and their correlation matrix ***R***. These data would be the input of our analysis.

Following the literature [61, 21], and particularly, the summary statistics version of SuSiE (SuSiE-RSS) [21], we have the following model of 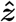,

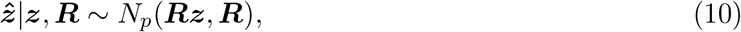

where ***z*** = (*z*_1_, *z*_2_, …, *z*_*p*_) denote the “standardized” true effect sizes. We use the same spike-and-slab prior for ***z***: when the variable *j* belongs to the group *k*,

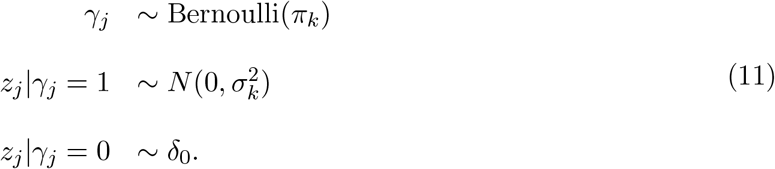

Again, we denote 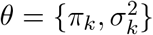 the prior parameters and *Γ* the causal configuration. The likeli-hood of this model is given by

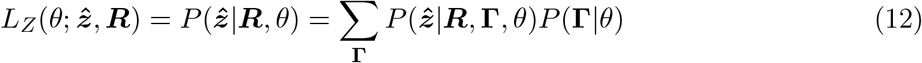

### Inference of the summary statistics model

We estimate the prior parameters *θ* by MLE. This can be done with the same algorithm used for the individual level model. Specifically, following SuSiE-RSS [21], the likelihood function under the individual level data can be rewritten in terms of the sufficient statistics 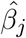 and 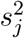. Then, if we make the following substitutions, the likelihood of the individual level model would be identical to that of the summary statistics model. Specifically, we change *β* = (*β*_1_, *β*_2_, …, *β*_*p*_) to ***z, X***^*T*^ ***X*** to ***R, X***^*T*^ ***y*** to 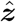, ***y***^*T*^ ***y*** to 1 and *n* to 1. Also the prior model of *z* in the summary statistics model is the same as the prior model of *β* in the individual level model. Therefore, we we can use the same EM algorithm and the update rules to estimate *θ*. The update rules follow Equations 5-6, where the PIP of variable *j* is now defined as 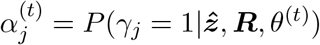 and the second moment of the posterior effect 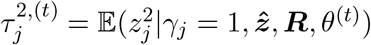.

Similarly as described in the inference procedures for individual level data, we divided genome into LD blocks, and used the blocks that are likely to have at most one effect, to infer the prior parameters *θ*. We then used these parameters to set the prior probabilities and prior variances of all variables within each block, and used SuSiE-RSS, with 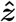, ***R*** as input and set *L* = 5, to obtain PIPs of all variables.

### Computation of marginal associations 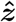 and correlation matrix *R*

When variable *j* is a gene, we can compute its Z-score, 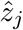 following the S-PrediXcan formula [31]. Specifically, suppose the gene *j* has *m* SNPs with nonzero weights in its expression prediction model. Let ***w***_*j*_ = (*w*_*j*1_, *w*_*j*2_, .., *w*_*jm*_) be their standardized weights, 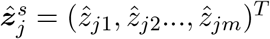 be the Z scores of the associations of the variants to the phenotype, and 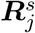 be the *m* × *m* correlation matrix, i.e. LD matrix, of these variants. Then 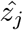 is basically the weighted sum of the Z-scores of all the *m* variants:

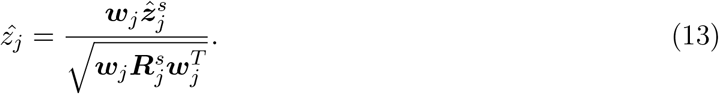

We also need ***R***, the correlation matrix, of all variables. We denote its element at row *i* and column *j* as *R*_*ij*_ = Cor(***X***_*i*_, ***X***_*j*_), where ***X***_*i*_ and ***X***_*j*_ are the *n*-dim. vectors (*n* is sample size) of the variables *i* and *j*. When both variable *i* and *j* are variants, *R*_*ij*_ is given by the variant LD matrix. When one of the variable, for example ***X***_*j*_, is an imputed gene, we can write it as: 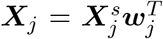, where 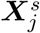 is a *n* × *m* matrix of genotypes for the *m* variants in gene *j* ^’^s prediction model. *R*_*ij*_ is then given by:

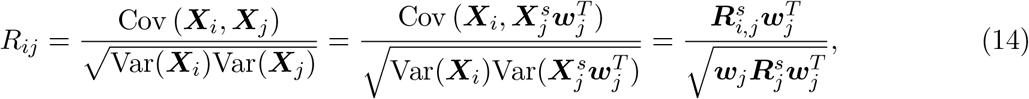

where 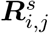 is the *m*-dimensional vector of correlation between variant *i* with each of the *m* variants in the prediction model of gene *j*. When both variables *i* and *j* are genes,

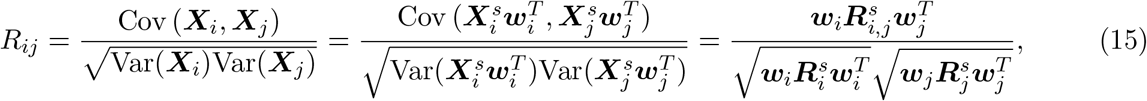

where 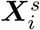 is the genotype matrix and ***w***_*i*_ is the vector of standardized weights for variants in gene *i*^’^s prediction model. 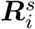 is the correlation matrix of 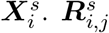 is now the correlation matrix between the variants in the prediction models of gene *i* and gene *j*.

### Estimating proportions of phenotypic variance explained by variants and genes

We assume that all the explanatory variables and the response variable in the regression model are standardized, with variance equal to 1. Then the proportion of variance explained (PVE) by a single variable *j*, is simply 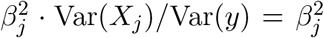. Assuming that we use the Z-scores in the summary statistical model, the effect size is related to Z-score by: 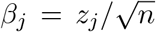, where *n* is the sample size. So on average, the PVE of a variable in group *k* (variant or gene) is: 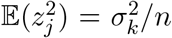, where *σ*_*k*_ is the prior variance of effect size in the group *k*, at the Z-score scale. The expected number of variables with non-zero effects in the group *k* is: *π*_*k*_ · |*M*_*k*_|, where *π*_*k*_ is the prior inclusion probability and |*M*_*k*_| the group size. Putting this together, the percent of total variance explained (PVE) by the group *k* is given by

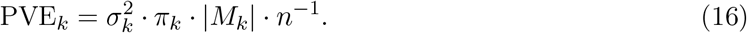

This equation is used to compute PVE from estimated parameters using both simulated and real data.

### Speeding up computation

Parameter estimation is the most time-consuming step. This is especially the case when variants are densely genotyped. We developed a “thinning” procedure to speed up this step. Specifically, we randomly select a fraction of SNPs in the single effect blocks, and only use these variants in the EM update. When most selected blocks have < 1000 variants, the EM algorithm can be done in one or two hours. We have found in our simulations that this thinning step does not affect the accuracy of results. In the final step of analyzing individual blocks, the estimated prior probability for variants from the thinning procedure will be scaled back for the original variants data. This is done by dividing the prior probability by the fraction of selected variants. The other parameters are not affected by this procedure. To reduce computational burden of the final step, we can analyze an individual block with the original variants only when results using the thinned variants showed strong signals for gene effect, e.g the maximum PIP of genes in the block > 0.8. Our software allows the user to specify the desired thinning parameter (the fraction of variants selected) in the parameter estimation step and the PIP threshold used to determine which blocks to analyze with original variants in the final step.

### Simulation procedure

In our simulations, we used the following data:

- Genotype data. We used genotype data from UK biobank by randomly selecting 80,000 samples. We then filtered samples to only keep “White British”, removed samples with missing information, mismatches between self-reported and genetic sex, or “outliers” as defined by UK Biobank. We also removed any individuals that have close relatives in the cohort. This ended up with a cohort of *n* = 45, 087 samples. We used SNPs from chr 1 to chr 22 and selected those with minor allele frequency > 0.05 and at least 95% genotyping rate. After filtering, 6,228,664 SNPs remained and were used in our analysis.
- Gene expression prediction models. We used GTEx v7 Adipose tissue dataset. This dataset contains 8,021 genes with expression models. We used the lasso weights from the FUSION website (http://gusevlab.org/projects/fusion/).

We first impute gene expression for all samples using the prediction models. SNP genotypes are harmonized between the expression prediction model and UK Biobank genotypes so that the reference and alternate alleles match. SNPs in the FUSION prediction models but not in UK Biobank, about 13% of all, were not used in imputing gene expression. We then sample the causal genes and SNPs under given prior inclusion probabilities *π*_*k*_’s, and then sample their effect sizes accordingly using the prior variance parameter 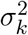. We then simulate *y* under the model defined in Equation 1. The prior parameters 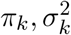 were chosen to reflect the genetic architecture in real data. In particular, it was estimated that gene expression mediates about 10-20% trait heritability [15]. And the studies using rare variants for complex traits suggested that about 5% of protein coding genes are likely causal [62]. Given these considerations, we set the prior probability for SNPs to 10^−4^ or 2.5 × 10^−4^, and PVE of SNPs to 0.3 or 0.5. For the genes, we set the prior probability 0.015 or 0.05 and PVE of genes from 0.02 to 0.1.

To test if our method is robust to mis-specified priors for causal gene effect, we have also simulated causal gene effect under the mixture of normal distributions. For the mixture of normal distributions, we used equal mixtures of four normal distributions, each with mean 0 and standard deviations with ratios of 1:2:4:8. That is for gene *j*, its prior distribution of causal effect size follows *β*_*j*_|*γ*_*j*_ = 1 *∼ ∑ ω ϵ[*`1,2,4,8]` ^π^ *′(N(0, ω σ ′*^*2*^). The prior probability being a casual gene is therefore 4*π ′* and causal effect size variance is 15*σ*′^2^. The prior probability of being a casual gene and the PVE of genes were set to values as described above.

To run cTWAS, we performed association of individual SNPs with the trait *y*, to obtain summary statistics of SNPs 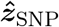. We randomly selected 2000 samples from the cohort to calculate SNP genotype correlation matrix, or LD matrix 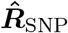. We then ran cTWAS summary statistics version under each simulation setting with 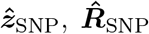 and expression prediction models as input. The software will harmonize SNP genotypes for 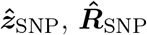 and expression prediction models, so that the reference and alternate allele match. To further reduce computational burden in estimating parameters, we only used 1 in every 10 SNPs (SNP thinning) in the EM algorithm. When calculating PIP, we first run SuSiE-RSS with *L* =5 in each LD Block with thinned SNPs. For each block with maximum gene PIP > 0.8, we re-run SuSiE-RSS with *L* = 5 with the original SNPs to get final gene PIPs.

### Running other methods on simulation data

We ran FUSION (https://github.com/gusevlab/fusion_twas) on the same simulated GWAS summary statistics data and the same gene prediction models. We ran coloc (https://cran.r-project.org/web/packages/coloc/index.html) which is provided as part of the FUSION package and used PIP4 > 0.8 as cutoff. We ran FOCUS (https://github.com/bogdanlab/focus) under the default setting and use PIP > 0.8 as the cut off. For SMR with HEIDI filter (https://yanglab.westlake.edu.cn/software/smr/), we used the significant ceQTL marginal association statistics provided by GTEx v7 as input, for genes with FUSION prediction models. These include significant variants ±1mb of the transcription start site (TSS) of each gene. For SMR, we used B-H adjusted *p* values < 0.05 to select significant genes, and we also required *p*_HEIDI > 0.05 for the HEIDI filter. For MR-JTI (https://github.com/gamazonlab/MR-JTI), we used the full eQTL marginal association statistics provided by GTEx v7 after LD-pruning (*r*^2^ = 0.2) as input. These include all variants ±1mb of the TSS of each gene. We analyzed genes with Fusion B-H adjusted *p* values < 0.05 using MR-JTI and required a Bonferroni-corrected *p* < 0.05 to determine significance. For PMR-Egger (https://github.com/yuanzhongshang/PMR), we used the full eQTL marginal association statistics provided by GTEx v7 as input, for genes with FUSION prediction models. We restricted the data to variants ±100kb of the body of each gene. Based on correspondence with the author of PMR-Egger, we regularized the LD matrices for these variants as 0.8***R*** + 0.2***I*** to avoid matrix decomposition errors. We used B-H adjusted *p* values < 0.05 to select significant genes for PMR-Egger.

### GWAS summary statistics

The LDL and SBP summary statistics were from the UK Biobank, computed by the Rapid GWAS project [29] using Hail [63]. These summary statistics were downloaded from the IEU OpenGWAS project [64] using GWAS IDs “ukb-d-30780_irnt” (LDL) and “ukb-a-360” (SBP). Both LDL and SBP summary statistics were based on the White British subpopulation of the UK Biobank, with sample sizes of N=343,621 and N=317,754 respectively. The IBD summary statistics were from the International IBD Genetics Consortium [65], computed by meta-analysis using METAL [66]. These summary statistics were obtained from IEU OpenGWAS using GWAS ID “ebi-a-GCST004131”. IBD includes cases of both Crohn’s disease and Ulcerative colitis. The IBD summary statistics were based on non-overlapping samples of European ancestry with a combined sample size of N=59,957. The SCZ summary statistics were from the Psychiatric Genetics Consortium and the CardiffCOGS study [67], computed by meta-analysis using METAL [66]. These summary statistics were obtained from the authors via the link provided in the manuscript. The SCZ summary statistics were based on non-overlapping samples of primarily European ancestry with a combined sample size of N=105,318.

### LD reference data

We computed the LD reference panel of common biallelic variants using the White British subpopulation of the UK Biobank. This panel is an in-sample reference for GWAS summary statistics from the Rapid GWAS project [29]. First, we used plate and well information from the genotyping to unambiguously identify over 99% (357,654 of 361,194) of the samples used in the Rapid GWAS project in our data. To ease computation, we randomly selected 10% of these samples to serve as the LD reference panel [68]. We also limited our panel to common autosomal variants with MAF > 0.01 in the Rapid GWAS analyses. Then, we computed correlations between all pairs of variants within each of 1,700 approximately independent regions. These regions are assumed to have low LD between them, and are based on previously identified regions [59] that could be lifted over from hg37 to hg38 positions. The final LD reference panel consists of 1700 correlation matrices and contains 9,309,375 variants. This LD reference panel was used when analyzing all traits, including those that were not measured in the White British subpopulation of UK Biobank.

### Prediction model of gene expression

We used expression prediction models for 49 GTEx [30] tissues from PredictDB [4] [31]. We used the mashr-based prediction models, which borrowed information across tissues to improve prediction accuracy in any specific tissue [41]. These prediction models were built using fine-mapped variants and are sparse, with a maximum of 5 eQTL per gene. The models were constructed using in-sample LD from GTEx, and the covariance between pairs of variants within each gene is reported alongside the models. We included only protein-coding genes for our analyses. Variants included in the models but missing in the GWAS summary statistics or LD reference were given zero weight.

### Harmonization of GWAS data and expression prediction models to LD reference

We restricted our analyses to variants that were non-missing in the GWAS summary statistics, expression prediction models, and LD reference panel. To ensure consistency between these three datasets, we performed two harmonization procedures. The objective of harmonization was to ensure that the reference and alternate alleles of each variant are defined consistently across all three datasets [21]. In our case, we must harmonize both the GWAS z-scores and the eQTL prediction models to our LD reference, and we use a different harmonize procedure for each. These procedures are based in part on previous work [31].

To describe the two procedures, it is necessary to define several cases of inconsistencies that can occur in either dataset. The first case is a variant with its reference and alternate alleles “flipped” with respect to the LD reference. The GWAS z-scores or eQTL weights in the prediction model of the flipped variants should have their signs reversed to be consistent with the LD reference. The second case is a variant that has had its strand “switched” with respect to the LD reference (e.g. variant is G/A in the LD reference but C/T in the other dataset). In this case, the reference and alternate alleles are the same, just named using different strands. The z-scores or weights of switched variants should not be changed, as they are already consistent with the LD reference. The third case is a variant that is “ambiguous” as to whether it is flipped or switched. This occurs when the two alleles of a variant are also complementary base pairs (A↔T substitutions or G↔C). For example, consider a variant that is A/T in the LD reference and T/A. It is unclear if this variant is flipped or switched with respect to the LD reference (both result in T/A), and it is ambiguous as to whether the signs of the z-scores or weights should be reversed. We say that variants are “unambiguous” when they do not involve substitutions of complementary base pairs.

To harmonize the z-scores from GWAS summary statistics, we first identified all inconsistencies in reference and alternate alleles between the z-scores and the LD reference. Next, we resolved all unambiguous cases of flipped and switched alleles, reversing the sign of z-scores that were flipped and taking no action for switched alleles. Then, we imputed the z-scores for ambiguous variants using all unambiguous variants in each of the LD regions [69]. If the sign of the imputed z-score did not match the sign of the observed z-score, we used the sign of the imputed z-score, reversing the sign of the observed z-score. Note that we did not perform the procedure to resolve ambiguous variants when analyzing LDL or SBP, as both the summary statistics and LD reference panel are derived from UK Biobank data.

To harmonize the eQTL prediction models, we first identified all inconsistencies in reference and alternate alleles between the prediction models and the LD reference. Next, we resolved all unambiguous cases of flipped and switched alleles, reversing the sign of weights that were flipped and taking no action for switched alleles. These steps to resolve unambiguous variants are the same as in the z-score harmonization procedure. To resolve ambiguous variants, we leveraged correlations between ambiguous and unambiguous variants in both our LD reference panel and the LD panel used to construct the PredictDB models. PredictDB reports the covariance between pairs of variants within each gene prediction model. For gene prediction models that include both ambiguous variants and unambiguous variants, we computed the sum of correlations between each ambiguous variant and the unambiguous variants in the prediction model, using both our LD reference panel and the LD used for the prediction models. If the sign of the total correlation in the LD reference of the prediction models did not match the sign of the total correlation in our LD reference panel, we reversed the sign of the prediction model weights for the ambiguous variant. If the total correlation in the LD reference was equal to zero, then we set the weight of the ambiguous variant to zero, as these ambiguous variants did not have any unambiguous variants in the same LD region. For gene prediction models that include only a single ambiguous variant and no unambiguous variants, we left the sign of the prediction model weight unchanged; the resulting gene z-score may have an incorrect sign, but the magnitude of the z-score will be correct. We excluded gene prediction models with multiple ambiguous variants and no unambiguous variants, as their gene z-scores could be incorrect in both sign and magnitude. Such exclusions were infrequent, affecting less than 1% of liver genes in the LDL analysis (94 of 11,502 genes with prediction models).

### Performing cTWAS analysis in real data

We used the following cTWAS settings when analyzing real data. For parameter estimation, we used the default procedure for selecting starting values of the E-M algorithm. We then performed 30 iterations of the E-M algorithm assuming L=1 effect (at most a single causal effect) in each region, using variants that were thinned by 10% to reduce computation. For computing PIPs of genes and variants, we used thinned variants and assumed L=5 (at most 5 causal effects) in each region. For regions with maximum gene PIP > 0.8, we recomputed PIPs using all variants, with L = 5. For this final step, we allowed a maximum of 20,000 variants in a region to reduce computation; if the maximum number of variants was exceeded, we randomly selected 20,000 variants to include. Unless specified otherwise, we used the threshold PIP > 0.8 for declaring significant genes.

### Total heritability and percentage of heritability mediated by genes using MESC

To assess parameter estimates from cTWAS, we computed total heritability and the percentage of heritability mediated by genes using MESC [15] for comparison. We used the same summary statistics as in the cTWAS analysis, and we used GTEx v8 expression scores for individual tissues available via the MESC repository. These expression scores are derived from the same GTEx dataset as the PredictDB models. Note that expression scores for “Kidney Cortex” tissue were not available from MESC, although this tissue is included in PredictDB and our other analyses.

### Silver standard and previously reported candidate genes of complex traits

To evaluate our findings, we compared the genes detected using cTWAS to curated lists of known or candidate genes for each trait. For LDL, we combined curated gene lists from two previous studies [13] [33], for a total of 69 LDL-related genes. These are genes that cause Mendelian disease or are drug targets, are in the KEGG “cholesterol metabolism” pathway, or are otherwise documented in the literature. For IBD, we used a list of 26 genes curated from literature, reported in the paper describing the activity-by-contact (ABC) model [70]. We refer to the gene lists for LDL and IBD as “silver standard” gene lists throughout the text based on the strength of their evidence. In supplemental materials, we also refer to “previously reported candidate genes” for SCZ and SBP. These are gene lists that are primarily supported by GWAS data rather than literature or experimental evidence. For SCZ, we used a list of 120 genes prioritized as SCZ-related based on proximity to GWAS, fine-mapping results and eQTL evidence [71]. For SBP, we used a list of 53 genes, supported by TWAS, gene expression, drug targets, pathway analysis and rare variant studies, reported in a large trans-ethnic GWAS (Table 3 of the paper) [72].

### Evaluating methods in distinguishing silver standard and bystander genes for LDL

Following earlier work [34], we assessed the performance of TWAS and cTWAS on real data by comparing their ability to distinguish LDL silver standard genes from other nearby genes. We defined a set of “bystander” genes that were within 1Mb of a silver standard gene. These bystandarder genes would be considered the negative set. We limited our analysis to the 46 of 69 silver standard genes with imputed expression after harmonization, and the 539 imputed bystander genes that are nearby these genes. Next, we determined if these silver standard and bystander genes were significant by TWAS (Bonferroni) or cTWAS (PIP > 0.8). Then, we computed the precision of each method, defined as: (# detected silver standard genes) / (# detected silver standard genes + # detected bystander genes).

### Fine-mapping analysis of POLK / HMGCR locus

To examine the POLK / HMGCR locus, we used fine-mapping results of LDL from UK Biobank from PolyFun-SuSiE [36] (downloaded from http://data.broadinstitute.org/alkesgroup/polyfun_results/biochemistry_LDLdirect.txt.gz). Specifically, PolyFun specifies prior probabilities for fine-mapping by estimating functional enrichments for a broad set of coding, conserved, regulatory and LD-related annotations. Then SuSiE was applied using the estimated priors with L=10 effects per locus.

To annotate the functions of variants their putative target genes, we used additional datasets. H3K4me1 ChIP-seq data of from human liver samples were downloaded from the ENCODE portal (ENCODE ID: “ENCFF323TFF”). Promoter Capture Hi-C (PC-HiC) data from human liver were obtained from a previous study [73]. Liver activity-by-contract (ABC) data were also obtained from a previous study [42].

### Classifying TWAS false positive genes for LDL by source of confounding

To better understand how TWAS generated false positive findings, we classified whether TWAS false positives as primarily due to confounding by variants or confounding by genes. We defined TWAS false positives as genes that were significant by TWAS (Bonferroni) but PIP < 0.5 by cTWAS. To categorize these false positive genes, we first assigned them to credible sets. These credible sets were reported by cTWAS, using the default SuSiE setting, which means that only credible sets with sufficient “purity” are reported (i.e. all variables in a credible set are highly correlated, r > 0.5). If a false positive gene was not included in any credible set, but was highly correlated (r > 0.5) with at least one variant or gene in a credible set, that false positive gene was also assigned to the credible set. After assigning a total of 83 false positive genes to credible sets, for each assigned gene, we summed the PIPs of all other genes and variants in its credible set to obtain total PIPs for confounding genes and variants. If the total gene PIP was higher than that of the variants, we classified the gene as confounded by genes, otherwise, confounded by variants.

### Assessing the novelty of cTWAS detected genes

If a gene was not included in the list of silver standard or previously reported genes of a trait, and was not the nearest gene of the lead variant in a genome-wide significant locus, we considered the gene a “novel” finding. We defined “nearest” genes as those with start or end positions that were the nearest out of all protein-coding genes to the lead variant of a genome-wide significant locus. To define independent lead variants for the GWAS, we performed the following procedure. First, we selected the most significant variant as a lead variant, and we removed all other variants within 500kb. Then, we iterated these selection and removal steps until there were no genome-wide significant variants remaining. We then identified the protein-coding genes that were nearest to each lead variant in the resulting list.

### Gene Ontology (GO) analysis of candidate genes

We performed enrichment analysis to characterize the genes detected by cTWAS for LDL and IBD. We used Enrichr [74] to identify Gene Ontology terms in the Biological Process, Molecular Function, or Cellular Component domains that are enriched for genes detected by cTWAS. Enrichr uses the Benjamini-Hochberg procedure to control FDR in each domain. We used a threshold of FDR < 0.05 for significant enrichment. We report the full enrichment analysis results using Enrichr across all domains for LDL and IBD (Table S3; Table S11). We compared the results from cTWAS genes with similar enrichment analysis results that used silver standard genes for LDL and IBD (Table S4; Table S12). We also compared these results with enrichment results from MAGMA [75] [76], a method that performs enrichment analysis based on GWAS data, using default settings (Table S5; Table S13). When reporting the IBD results (Table S9), we annotated detected genes with the enriched GO terms (using cTWAS genes, silver standard genes, or MAGMA) that each gene is a member of. For all three GO annotations, a maximum of 5 significant terms per gene were shown, ordered by odds ratios (cTWAS, silver standard) or p-values (MAGMA).

To visualize the GO biological process terms enriched for LDL cTWAS genes (Fig. 5B), we removed redundant terms to improve clarity. To do this, we first ranked all GO terms by their significance (p-values). A term was considered redundant if all the cTWAS genes in that term had already been included in a more significant GO term.

To visualize the GO biological process terms enriched for IBD cTWAS genes (Fig. 6E), we used the “Weighted Set Cover” tool in WebGestalt [43] to remove redundant GO terms. To further simplify visualization, we omitted GO terms whose detected genes were all included in other terms identified by Weighted Set Cover. We also report the full enrichment analysis results using WebGestalt for IBD (Table S10).

### Summarizing cTWAS results using tissue groups

To aid the interpretation of cTWAS findings, we grouped related tissues into “tissue groups” and summarized the findings within these groups. We used previously defined tissue groups that assigned 37 of the 49 tissues to one of 7 tissue groups [15]. We then took the union of genes detected at PIP > 0.8 in any tissue within each tissue group, and we used these combined lists of detected genes for downstream analyses.

