## Supplementary_notes for "Adjusting for genetic confounders in transcriptome-wide association studies leads to reliable detection of causal genes"

#### EM algorithm for estimating the prior parameters

We first restate our full model as described in Methods. We have a linear model with $K$ groups of explanatory variables:

$$y = \sum_{k=1}^{K} \sum_{j \in M_k} \beta_j X_j + \epsilon, \qquad \epsilon \sim N(0, \sigma^2), \tag{1}$$

where $X_j, 1 \leq j \leq p$, is $j$-th explanatory variable and $j \in M_k$ denotes that it belongs to the group $k$. The effect size $\beta_j$'s follow spike-and-slab prior distributions with group-specific parameters. Let $\gamma_j$ be an indicator of whether $X_j$ has non-zero effect:

$$\begin{array}{ll} \gamma_j & \sim \text{Bernoulli}(\pi_k) \\ \beta_j | \gamma_j = 1 & \sim N(0, \sigma_k^2) \\ \beta_j | \gamma_j = 0 & \sim \delta_0. \end{array} \tag{2}$$

For simplicity, we assume $\sigma^2$ is given (see Methods). Our goal is to estimate the prior parameters $\theta = \{\pi_k, \sigma_k^2, 1 \leq k \leq K\}$.

We use Expectation-Maximization (EM) algorithm to estimate these parameters, treating the effect sizes $\boldsymbol{\beta} = (\beta_1, \cdots, \beta_p)$ and the configuration of all indicators $\boldsymbol{\Gamma} = (\gamma_1, \cdots, \gamma_p)$ as missing data. The complete data log-likelihood function is given by:

$$\begin{aligned} \log P(\boldsymbol{y}, \boldsymbol{\Gamma}, \boldsymbol{\beta} | \boldsymbol{X}, \theta) &= \log P(\boldsymbol{\Gamma}|\theta) + \log P(\boldsymbol{y}, \boldsymbol{\beta} | \boldsymbol{X}, \boldsymbol{\Gamma}, \theta) & (3) \\ &= \log P(\boldsymbol{\Gamma}|\theta) + \log P(\boldsymbol{\beta}|\boldsymbol{\Gamma}, \theta) + \log P(\boldsymbol{y}|\boldsymbol{X}, \boldsymbol{\beta}) & (4) \\ &= \sum_{k=1}^{K} \sum_{j \in M_k} \log P(\gamma_j|\pi_k) + \sum_{k=1}^{K} \sum_{j \in M_k} \log P(\beta_j|\gamma_j, \sigma_k^2) + \log P(\boldsymbol{y}|\boldsymbol{X}, \boldsymbol{\beta}). & (5) \end{aligned}$$

Note that the last term does not depend on the prior parameters $\theta$, so we can ignore it in the EM algorithm.

In the E-step, we take expectation of the complete log-likelihood function over $\boldsymbol{\beta}, \boldsymbol{\Gamma} | \theta^{(t)}$, where $\theta^{(t)}$ is the parameters from step $t$:

$$\begin{aligned} Q(\theta|\theta^{(t)}) &= \mathbb{E}_{\boldsymbol{\beta}, \boldsymbol{\Gamma}} \log P(\boldsymbol{y}, \boldsymbol{\Gamma}, \boldsymbol{\beta} | \boldsymbol{X}, \theta, \sigma) & (6) \\ &= \sum_{k=1}^{K} \sum_{j \in M_k} \mathbb{E}_{\gamma_j} \log P(\gamma_j|\pi_k) + \sum_{k=1}^{K} \sum_{j \in M_k} \mathbb{E}_{\beta_j, \gamma_j} \log P(\beta_j|\gamma_j, \sigma_k^2) + \text{constant}. & (7) \end{aligned}$$

We can simplify this function. First, we note:

$$\mathbb{E}_{\gamma_j} \log P(\gamma_j|\pi_k) = \mathbb{E}_{\gamma_j}[\gamma_j \log \pi_k + (1 - \gamma_j) \log(1 - \pi_k)] = \alpha_j^{(t)} \log \pi_k + (1 - \alpha_j^{(t)}) \log(1 - \pi_k), \tag{8}$$

where $\alpha_j^{(t)} = P(\gamma_j = 1 | \theta^{(t)}, \boldsymbol{X}, \boldsymbol{y})$ is the posterior inclusion probability of variable $j$ at the $t$-th iteration. Next,

$$\begin{aligned} \mathbb{E}_{\beta_j, \gamma_j} \log P(\beta_j|\gamma_j, \sigma_k^2)) &= \mathbb{E}_{\beta_j, \gamma_j}[\gamma_j \log P(\beta_j|\gamma_j = 1, \sigma_k^2) + (1 - \gamma_j) \log P(\beta_j|\gamma_j = 0)] & (9) \\ &= \alpha_j^{(t)} \cdot \mathbb{E}_{\beta_j|\gamma_j=1}[\log P(\beta_j|\gamma_j = 1, \sigma_k^2)] + (1 - \alpha_j^{(t)}) \cdot \mathbb{E}_{\beta_j|\gamma_j=0}[\log P(\beta_j|\gamma_j = 0)]. & (10) \end{aligned}$$

Given that $\beta_j = 0$ when $\gamma_j = 0$, the second term in the expectation is simply 0. For the first term, we plug in the normal density $\beta_j \sim N(0, \sigma_k^2)$:

$$\mathbb{E}_{\beta_j,\gamma_j} \log P(\beta_j|\gamma_j, \sigma_k^2)) = \alpha_j^{(t)} \cdot \mathbb{E}_{\beta_j|\gamma_j=1} \left( -\log \sqrt{2\pi}\sigma_k - \frac{\beta_j^2}{2\sigma_k^2} \right) = \alpha_j^{(t)} \cdot \left( -\log \sqrt{2\pi}\sigma_k - \frac{\tau_j^{2,(t)}}{2\sigma_k^2} \right), \tag{11}$$

where $\tau_j^{2,(t)} = \mathbb{E}(\beta_j^2|\gamma_j = 1, \boldsymbol{X}, \boldsymbol{y}, \theta^{(t)})$ is the second moment of posterior effect size distribution of $\beta_j$ given the current parameter estimate $\theta^t$ and that $\gamma_j = 1$.

Putting all these together, we have:

$$Q(\theta|\theta^{(t)}) = \sum_{k=1}^{K} \sum_{j \in M_k} \left[ \alpha_j^{(t)} \log \pi_k + (1 - \alpha_j^{(t)}) \log(1 - \pi_k) \right] + \sum_{k=1}^{K} \sum_{j \in M_k} \alpha_j^{(t)} \left( -\log \sqrt{2\pi}\sigma_k - \frac{\tau_j^{2,(t)}}{2\sigma_k^2} \right) \tag{12}$$

At the M-step, we find $\theta$ that maximize $Q(\theta|\theta^t)$. We begin with the update rule for $\pi_k$, noting that only the first term contains $\pi_k$.

$$\frac{\partial Q(\theta|\theta^{(t)})}{\partial \pi_k} = \sum_{j \in M_k} \left[ \frac{\alpha_j^{(t)}}{\pi_k} - \frac{1 - \alpha_j^{(t)}}{1 - \pi_k} \right] = 0. \tag{13}$$

Solving this equation gives the update rule of $\pi_k$:

$$\pi_k^{(t+1)} = \frac{1}{|M_k|} \sum_{j \in M_k} \alpha_j^{(t)}, \tag{14}$$

where $|M_k|$ is the size of $M_k$. Next we derive the update rule of $\sigma_k^2$:

$$\frac{\partial Q(\theta|\theta^{(t)})}{\partial \sigma_k} = \sum_{j \in M_k} \alpha_j^{(t)} \left[ -\frac{1}{\sigma_k} + \frac{\tau_j^{2,(t)}}{\sigma_k^3} \right] = 0. \tag{15}$$

This leads to the update rule for $\sigma_k^2$ as:

$$\sigma_k^{2,(t+1)} = \frac{\sum_{j \in M_k} \alpha_j^{(t)} \tau_j^{2,(t)}}{\sum_{j \in M_k} \alpha_j^{(t)}}. \tag{16}$$

The computation of $\alpha_j^{(t)}$ and $\tau_j^{2,(t)}$ at each iteration will be described below. We perform E and M step until each prior parameters change $< 0.00001$ in one iteration or the maximum number of iterations is met.

### Calculation of $\alpha_j$ and $\tau_j^2$ under the single effect approximation.

In the EM iteration, we need to compute the PIP $\alpha_j^{(t)}$ and the posterior second moment $\tau_j^{2,(t)}$ of each variable (the index $(t)$ will be dropped later for simplicity). To simplify, we analyze each block one at a time, assuming each block has at most a single variable with non-zero effects. We further assume that each single block explains a minimal variance of $\boldsymbol{y}$, so we set $\sigma = 1$ in the calculation. Let $\Gamma_j$ be the configuration where $\gamma_j = 1$ and all other indicators are 0, and $\boldsymbol{\Gamma}_0$ be the null configuration where $\gamma_j = 0$ for all variables. Let $p_j$ be the prior inclusion probability of variable $j$, which depends on the group variable $j$ belongs to, $p_j = \pi_k$ when $j \in M_k$. The prior probabilities of the configurations are given by:

$$P(\boldsymbol{\Gamma}_0|\theta) = \prod_j (1 - p_j), \tag{17}$$

$$P(\boldsymbol{\Gamma}_j|\theta) = \frac{p_j}{1 - p_j} \cdot P(\boldsymbol{\Gamma}_0|\theta). \tag{18}$$

For variable $j$, we define its Bayes factor, as the probability of data under $\mathbf{\Gamma}_j$ vs. under $\mathbf{\Gamma}_0$. Based on the Wakefield formula[1], it is given by:

$$\begin{aligned} B_j &:= \frac{P(\boldsymbol{y}|\boldsymbol{X}, \mathbf{\Gamma}_j, \theta)}{P(\boldsymbol{y}|\boldsymbol{X}, \mathbf{\Gamma}_0, \theta)} && (19) \\ &= \sqrt{\frac{s_j^2}{\sigma_k^2 + s_j^2}} \cdot \exp\left(\frac{\hat{\beta_j}^2 \sigma_k^2}{2 s_j^2 (\sigma_k^2 + s_j^2)}\right), && (20) \end{aligned}$$

where $\sigma_k$ is the prior variance of effect size for variables in group $k$. $\hat{\beta}_j$ is the estimated effect size of variable $j$, and $s_j$ is the standard error. They are given by:

$$\begin{aligned} \hat{\beta}_j &:= \boldsymbol{X}_j^T \boldsymbol{y}/n, && (21) \\ s_j^2 &:= (\boldsymbol{y} - \boldsymbol{X}_j \hat{\beta}_j)^T (\boldsymbol{y} - \boldsymbol{X}_j \hat{\beta}_j)/(n \boldsymbol{X}_j^T \boldsymbol{X}_j) \approx 1/n. && (22) \end{aligned}$$

The posterior inclusion probabilities depend on the prior and the BFs of variables:

$$\begin{aligned} \alpha_j &= P(\gamma_j = 1|\boldsymbol{X}, \boldsymbol{y}, \theta) && (23) \\ &= P(\mathbf{\Gamma}_j|\boldsymbol{X}, \boldsymbol{y}, \theta) && (24) \\ &= \frac{P(\boldsymbol{y}|\boldsymbol{X}, \mathbf{\Gamma}_j, \theta) P(\mathbf{\Gamma}_j|\theta)}{P(\boldsymbol{y}|\boldsymbol{X}, \mathbf{\Gamma}_0, \theta) P(\mathbf{\Gamma}_0|\theta) + \sum_j P(\boldsymbol{y}|\boldsymbol{X}, \gamma_j, \theta) P(\mathbf{\Gamma}_j|\theta)} && (25) \\ &= \frac{P(\mathbf{\Gamma}_j|\theta) \cdot B_j}{P(\mathbf{\Gamma}_0|\theta) + \sum_j P(\mathbf{\Gamma}_j|\theta) \cdot B_j} && (26) \\ &= \frac{p_j/(1 - p_j) B_j}{1 + \sum_j p_j/(1 - p_j) B_j}. && (27) \end{aligned}$$

Following the derivation in the SuSiE paper, the posterior distribution of $\beta_j|\gamma_j = 1$ follows $N(\mu_{1j}, \sigma_{1j}^2)$ where $\sigma_{1j}^2 = s_j^2 \sigma_k^2/(\sigma_k^2 + s_j^2)$ and $\mu_{1j} = \sigma_{1j}^2 \hat{\beta}_j / s_j^2$, so we have the second moment of posterior distribution,

$$\tau_j^2 = \mathbb{E}(\beta_j^2|\gamma_j = 1, \theta) = \text{var}(\beta_j|\boldsymbol{X}, \boldsymbol{y}, \theta, \gamma_j = 1) + (\mathbb{E}(\beta_j|\boldsymbol{X}, \boldsymbol{y}, \theta, \gamma_j = 1))^2 = \mu_{1j}^2 + \sigma_{1j}^2. \tag{28}$$

When using summary statistics in the form of Z-scores, we replace $\hat{\beta}_j$ as $\hat{z}_j/n$, where $\hat{z}_j$ is the Z-score of variable $j$. $\alpha_j$ can still be derived by Equation 27 where $B_j$ is given by

$$B_j = \sqrt{\frac{s_j^2}{\sigma_k^2 + s_j^2}} \cdot \exp\left(\frac{\hat{z_j}^2 \sigma_k^2}{2 s_j^2 (\sigma_k^2 + s_j^2)}\right). \tag{29}$$

where $\sigma_k$ is the variance of Z score for variables in group $k$. $\tau_j^2$ can still be derived by Equation 28 where $\mu_{1j} = \sigma_{1j}^2 \hat{z}_j$.

In the implementation of our method, we used the algorithm of single effect model (SER) from SuSiE to get $\alpha_j$ and $\tau_j^2$. When $p_j \ll 1$, PIP and the second moment of posterior distribution under SuSiE SER should be similar or the same as those from our model. In more detail, we use $p_j$ as prior probabilities in SuSiE and $\pi_0 = 1 - \sum_j p_j$ as null weight, PIP under SuSiE SER $\alpha'_j$ is given by

$$\alpha'_j = \frac{p_j B_j}{\pi_0 + \sum_j p_j B_j} = \frac{p_j/\pi_0 \cdot B_j}{1 + \sum_j p_j/\pi_0 \cdot B_j} \approx \alpha_j. \tag{30}$$

For the approximation, we used $\pi_0 = 1 - \sum_j p_j \approx 1 - p_j$ under the assumption that $p_j \ll 1$. The posterior distribution of $\beta_j|\gamma_j = 1$ under SuSiE SER is the same as in our model.

Similarly, we used the algorithm of the SER model in SuSiE RSS to get $\alpha_j$ and $\tau_j^2$ when the input is summary statistics($\hat{\boldsymbol{z}}, \boldsymbol{R}$).

### Initialization and selection of single effect blocks

To have a good chance of working with blocks satisfying the single effect assumption, we compute the prior probability of a block having at most one causal variable, denoted as $p_{\text{single effect}}$. It is given by:

$$p_{\text{single effect}} = P(\mathbf{\Gamma}_0) + \sum_j P(\gamma_j) = \prod_j (1 - p_j) \left( 1 + \sum_j \frac{p_j}{1 - p_j} \right). \quad (31)$$

Note that this probability considers both cases where a block has no causal variable, and just one. We choose blocks with $p_{\text{single effect}}$ above a threshold, 0.8 in our simulation and real data analysis, to be used in prior parameter estimation. The effect of this block selection step is that large blocks that are likely to contain more than one signal *a priori*, are filtered.

To initialize the EM algorithm described in "Inference of the individual level model" and "Inference of the summary statistics model", we use the same prior parameters for genes and variants. The thinning procedure is applied at the beginning of the EM algorithm if specified. We then run 3 iterations of the EM algorithm using all LD blocks, using this as an initial estimate of $p_j$. Then, we select single effect blocks based on $p_{\text{single effect}}$ given by Equation (31) and used these for 30 more iterations of the EM algorithm.

### Connection with colocalization methods

We show that under certain conditions, cTWAS reduces to colocalization analysis. Specifically, we assume there is a single causal gene in a region being studied, and that gene has a single causal eQTL variant (known as eQTN in literature). Let $X$ be the expression of the causal gene, $\tilde{X}$ be its *cis*-genetic component, and $G_j$ be the genotype of the eQTN. We have:

$$\tilde{X} = G_j w_j, \quad (32)$$

where $w_j$ is the weight of $G_j$ in the prediction model of $X$. We also assume that $G_j$ acts on the phenotype only through the gene $X$, thus its direct effect $\theta_j = 0$. Our model of the phenotype $y$ is given by:

$$y = \tilde{X}\beta + \sum_{i \neq j} G_i \theta_i + \epsilon, \quad (33)$$

where $\beta, \theta_i$ are the causal effects of the gene and variants, respectively. Also note that both $\beta$ and $\theta_i$'s follow spike-and-slab prior with prior inclusion probabilities $\pi_G$ (gene effect) and $\pi_V$ (variant effect), respectively. We can now plug in $\tilde{X} = G_j w_j$ into the model of phenotype:

$$y = G_j w_j \beta + \sum_{i \neq j} G_i \theta_i + \epsilon. \quad (34)$$

We see now that the gene effect $\beta \neq 0$ if and only if the effect of variant $j$, $w_j \beta \neq 0$. So our problem reduces to a fine-mapping problem, where we determine which variant(s) has non-zero coefficients. The difference with standard fine-mapping is that the prior inclusion probabilities vary with variants. The eQTN variant has prior probability $\pi_G$, while other variants have prior $\pi_V$, generally smaller than $\pi_G$. So the model would effectively perform fine-mapping, favoring the eQTN variant.

The model we described is effectively Enloc, a method for colocalization analysis [**wenintegrating**]. We give a short description here of Enloc here. We denote $d_j$ as an indicator of whether $G_j$ is an eQTL, and $\gamma_j$ as an indicator of whether $G_j$ is a causal variant of the phenotype in GWAS. Colocalization of eQTL and GWAS trait in a region of interest thus means that there is some variant in the region, where $d_j = 1$ and $\gamma_j = 1$. Enloc performs colocalization analysis in several steps. It first performs fine-mapping analysis of eQTL data, estimating $P(d_j = 1)$ for all variants. In the next step, Enloc assesses how often the eQTNs are also causal variants of a complex phenotype from GWAS. This step estimates the conditional distribution $P(\gamma_j | d_j)$. In general, we expect $P(\gamma_j = 1 | d_j = 1) >> P(\gamma_j = 1 | d_j = 0)$. The enrichment of causal variants in eQTNs is reflected by the parameter $\alpha_1$ in Enloc, defined as the log odds ratio:

$$\alpha_1 = \log \frac{P(\gamma_j = 1|d_j = 1)/P(\gamma_j = 0|d_j = 1)}{P(\gamma_j = 1|d_j = 0)/P(\gamma_j = 0|d_j = 0)}. \tag{35}$$

Enloc uses the data across genome to estimate $\alpha_1$. In the final step, Enloc performs fine-mapping of the GWAS trait in any regions of interest, using the prior probabilities of $\gamma_j$'s estimated in the previous step. Because eQTNs generally have higher prior, this allows Enloc to favor eQTNs in fine-mapping of the trait GWAS. The results of Enloc are expressed as $P(d_j = 1, \gamma_j = 1|D, \hat{\alpha_1})$, where $D$ means all the data we have.

From this description of Enloc, we can see that the reduced cTWAS model, Equation 34, is almost equivalent to Enloc. Both methods perform fine-mapping of causal variants of a phenotype, favoring eQTNs. In fact, the prior probabilities of the two models are related by:

$$P(\gamma_j = 1|d_j = 0) \approx \pi_V, \qquad P(\gamma_j = 1|d_j = 1) = \pi_G. \tag{36}$$

To see this, we note that under cTWAS, the prior probability of a non-eQTN variant being causal variant to the GWAS trait is $\pi_V$. The prior probability of an eQTN being causal variant is just the prior probability that its target gene is causal, which is $\pi_G$. We can now express $\alpha_1$, the key Enloc parameter, in terms of $\pi_G$ and $\pi_V$:

$$\alpha_1 = \log \frac{\pi_G/(1-\pi_G)}{\pi_V/(1-\pi_V)} \approx \log \frac{\pi_G}{\pi_V}, \tag{37}$$

where we use the approximation that $\pi_G, \pi_V$ are usually small, so $1 - \pi_G \approx 1$ and $1 - \pi_V \approx 1$. There is one subtle difference between this reduced cTWAS model and Enloc. Enloc uses a common parameter for the prior variance of effect sizes for all variants, including eQTNs. cTWAS in contrast, allows the prior variance to differ between gene effects (hence eQTL effects) and variant effects. In our simulations, we have found that using a common prior variance for gene and variant effects leads to inflated PIPs (data not shown).

Having shown the near equivalence of Enloc and the reduced cTWAS model, we can see that cTWAS is also related to coloc. coloc performs model selection, comparing several models, $H_1$: there is an eQTL in the region being analyzed but no causal variant for GWAS, $H_2$: there is a causal variant for GWAS trait but no eQTL, $H_3$: there are distinct eQTL and GWAS causal variants, and $H_4$: the same causal variant affects both expression and phenotype. The goal of coloc is largely about estimating the posterior probability of $H_4$, i.e. $P(H_4|D)$, where $D$ denotes the eQTL and GWAS data we have. This probability is related to the Enloc model by:

$$P(H_4|D) = \sum_j P(d_j = 1, \gamma_j = 1|D). \tag{38}$$

Having shown the connection between coloc and Enloc, we can also relate the key parameters under the two models. In coloc, the prior probabilities of the four models are related to the prior inclusion probabilities of variants: $p_1$, the prior of a variant being eQTN; $p_2$, the prior of causal variant of GWAS trait; and $p_{12}$, the prior of a variant being causal to both expression and GWAS trait. Whether coloc detects a colocalization, i.e. $H_4$ is chosen, is controlled largely by these prior parameters, $p_1, p_2, p_{12}$. In the Enloc paper, the authors show that Enloc is equivalent to coloc with the parameters under the two methods related by (see Equation 10 of Enloc):

$$\alpha_1 = \log \frac{p_{12}(1 - p_1 - p_2 - p_{12})}{p_1 p_2} \approx \log \frac{p_{12}}{p_1 p_2}, \tag{39}$$

where we again assume small prior probabilities. We can also see that the coloc parameteres are related to the cTWAS parameters. In fact, $p_1$ is the prior of causal variant, so $p_1 \approx \pi_V$; and $p_{12}/p_2$ is the conditional probability that a variant is causal to the phenotype given that it is an eQTN. As we have explained above, this is $P(\gamma_j = 1|d_j = 1)$, which is $\pi_G$ under the reduced cTWAS model. So the enrichment parameter under coloc becomes:

$$\log \frac{p_{12}}{p_1 p_2} = \log \frac{p_{12}/p_2}{p_1} \approx \log \frac{\pi_G}{\pi_V}. \tag{40}$$

This is the same Equation 37 we have derived linking $\alpha_1$ in Enloc and prior parameters of cTWAS.

From these derivations, we see that under the simple scenario of single causal gene, and single eQTN, cTWAS is equivalent to Enloc and coloc. Compared to coloc, both Enloc and the reduced cTWAS have the advantage that all the key prior parameters are estimated from the genome-wide data analysis. In contrast, coloc sets the parameters, $p_1, p_2, p_{12}$, somewhat arbitrarily. Many have reported that coloc results are sensitive to these parameters. Of course, the equivalence of cTWAS with colocalization methods only applies in the simple scenario. In general, cTWAS has the advantage that it allows multiple causal genes, and multiple eQTNs.

### Detailed simulation results for PMR-Egger

While all competing methods suffered from high false positive rates in our simulations, PMR-Egger performed particularly poorly. Its false rate rate was even higher than standard TWAS (FUSION). This is quite unexpected, given that the objective of PMR-Egger is to avoid false positives under TWAS. We thus investigated the simulation results for PMR-Egger in more detail to understand the reasons for its poor performance.

We begin with a short review of PMR-Egger. PMR-Egger analyzes one gene a time. Let $X$ be the expression of a gene of interest, and $\tilde{X}$ be its *cis*-genetic component. PMR-Egger assumes that the trait $y$, depends on $\tilde{X}$, as well as the pleiotropic effect of nearby variants, whose genotypes are denoted as $Z_y$ ($p$-dimension). We have:

$$y = \mu_y + \tilde{X}\alpha + Z_y\gamma + \epsilon_y, \tag{41}$$

where $\mu_y$ is the mean trait value, $\alpha$ is the gene effect, and $\gamma$ the $p$-dimensional effect sizes of nearby variants. PMR-Egger also accounts for the uncertainty of $\tilde{X}$ through another model linking observed expression $X$ with genotypes of all nearby variants under a polygenic assumption. The model as shown in Equation 41, by itself, is not identifiable, because of colinearity of $\tilde{X}$ and $Z_y$: $\tilde{X}$ is effectively a linear combination of variant genotypes. Thus PMR-Egger made an extra assumption, known as Egger regression in Mendelian Randomization, that all variants have identical pleiotropic effects: $\gamma_1 = \gamma_2 = \cdots = \gamma_p = \gamma$. The goal of PMR-Egger is to test if $\alpha = 0$, while allowing $\gamma \neq 0$.

When we first ran PMR-Egger with default settings, we noted that it did not return results for a large percentage of genes (83.4% on average for the high PVE simulations) due to errors decomposing LD matrices. Tuning the shrinkage parameter ($\lambda$), as suggested by the software, did not substantially reduce the number of decomposition errors. Upon discussion with the author, we performed an alternative regularization of the LD matrices, regularizing them as $0.8\boldsymbol{R} + 0.2\boldsymbol{I}$, where $\boldsymbol{R}$ is the original LD matrix, and $\boldsymbol{I}$ the identity matrix. The number of decomposition errors was substantially reduced, although a substantial fraction of genes still have errors (18.6% on average for the high PVE simulations).

Among the genes that were significant by PMR-Egger, the proportion of false positives was very high (85.9% on average for the high PVE simulations), higher than FUSION. To understand this discrepancy, we focused on a single simulation from the high PVE scenario and the 6,454 genes which had both FUSION and PMR-Egger results. We observed that $-\log_{10} p$-values for FUSION and PMR-Egger are generally correlated ($r = 0.567$), but there are outlier genes which are insignificant by FUSION but highly significant by PMR-Egger (Fig. S13A). Many of these outlier genes are confounded by nearby causal variants with relatively large effect sizes.

As an example, we visualized the locus containing the gene *ICA1*, which was insignificant by FUSION, but had very significant $p$-values ($\sim 10^{-29}$) by PMR-Egger (Fig. S13B). At this locus, there is a single causal variant with a large effect size, and no causal gene. As expected, the causal variant shows the strongest association in the locus (Fig. S13B, top, the red vertical bar). The FUSION expression prediction model of *ICA1* has two variants, both of which are nearby (within 72kb) the causal variant (Fig. S13B, middle), but are not in LD with the causal variant ($R^2 = -0.022$ for strongest correlation). In the eQTL analysis of *ICA1*, the two variants in the prediction model show strong associations, as expected, but the causal variant is also nominally significant (Fig. S13B, bottom). Because PMR-Egger uses all *cis*-variants of a gene to predict expression under a polygenic assumption, it is likely that the prediction model of PMR-Egger includes some contribution of the causal variant. This induces a correlation of predicted expression of *ICA1* with the simulated trait. On the other hand, the null model of PMR-Egger, where $\alpha = 0$, attempts to explain the data using an identical effect for all *cis*-variants of *ICA1*. However, this is a mismatch of the data, as the association is simply due to a single causal variant with large effect. Taken together, we believe these two facts (the correlation of $\tilde{x}$ and $y$ due to PMR-Egger's expression prediction model, and the poor fit of the data by the null model) explain the false positive finding by PMR-Egger in this case. The FUSION method, on the other hand, avoided this mistake. The two variants of *ICA1* are not in high LD with the causal variant, and have low associations with the trait in GWAS (Fig. S13B, top). As a result, the imputed expression using FUSION is not highly correlated with the trait.

Many other genes that are discordant between FUSION and PMR-Egger are qualitatively similar. These results suggest that PMR-Egger is susceptible to false positives, when (1) the underlying expression prediction model is sparse with only a few causal variants, and (2) when there is a nearby large effect causal variant acting on the phenotype directly.
