## Supplementary_figures_tables_legend for "Adjusting for genetic confounders in transcriptome-wide association studies leads to reliable detection of causal genes"

### Supplemental Figures

**Figure S1. Causal diagrams representing several scenarios of how genetic variant, expression and phenotype are related.** $X$ and $X'$, gene expression; $G$ and $G'$ are genotypes of variants affecting expression of the two genes; $\tilde{X}$ and $\widetilde{X'}$,, *cis*-genetic component of gene expression (*i.e.* imputed expression); $y$, phenotypic trait; $U$, unmeasured confounder. Arrows indicate causal effect; double-headed arrows between $G$ and $G'$ indicate LD. Note that it is possible that $G$ and $G'$ are identical variants. This happens when the same variant affects both $X$ and $X'$,. **A.** Reverse causality: the phenotype $y$ has an effect on the expression of gene $X$. There are two paths from $X$ to $y$, either through $U$ or not. But in both paths, $X$ is a collider that blocks the association of $X$ and $y$. **B.** Confounding by a nearby gene $X'$ creates correlation of $X$ and $y$. The LD between eQTLs of the two genes creates a non-causal, backdoor path (shown in red) between $X$ and $y$. **C.** Confounding by a nearby variant $G'$ creates correlation of $X$ and $y$. The backdoor path from $X$ to $y$ through $G'$ is shown in red.

**Figure S2. Additional simulation results. A.** Parameter estimation results for variant effect related parameters. From left to right, percentage of causal variants ($\pi_V$), standard deviation of causal variants effect size and PVE from variants. The high and low PVE settings are the same as described in Figure 2. **B.** Gene PIP calibration plots under additional settings. For each plot, we listed the parameters used in simulating the trait at the top. $\pi_G$ is prior probability for a gene being causal, $\mathrm{PVE_G}$ is PVE by gene effect; $\pi_V$ is prior probability for a variant being causal, $\mathrm{PVE_V}$ is PVE by variant effect. Under each setting, gene PIPs from 5 simulations are grouped into bins. The plot shows the proportion of true causal genes (Y axis) against the average PIPs (X axis) under each bin. A well calibrated method should produce points along the diagonal lines (red). +/- standard error is shown for each point in the vertical bars.

**Figure S3. Results from simulations using mixtures of normal distribution as the prior distribution of effect sizes.** From A to J, each panel shows results from one simulation setting, as represented by true parameter values, shown as horizontal bars in the panels. The first 4 figures in each panel show parameter estimation results. $G$ is the prior probability for a gene being causal; $\pi_V$ is the prior probability for a SNP being causal; enrichment is defined as $\pi_G/\pi_V$; gene PVE and variant PVE are the percent of phenotypic variance explained by genes and variants, respectively. Each dot represents the result from one out of five simulations. Horizontal bars show the true parameter values. The last figure under each simulation setting is the PIP calibration plot, similarly as described in Figure 2B and Figure S2B.

**Figure S4. Additional results for cTWAS analysis of LDL cholesterol using liver eQTLs.**
**A.** Convergence of estimated parameters over 30 iterations of the E-M algorithm. "Proportion Causal" is the prior inclusion probability of genes and variants. "Effect Size" is the prior variance of the gene and variant effect sizes. "Enrichment" is the ratio of gene to variant prior inclusion probabilities. "PVE" is the proportion of variance explained by genes and variants. **B.** cTWAS results at the *PRKD2* locus. Figure legend is the same as in the locus plots in the main figures, see Fig. 4B.

**Figure S5. Mean number of imputed genes across all tissues for IBD, SCZ and SBP.** The numbers of imputed genes analyzed by cTWAS vary slightly among the three traits, because of the differences in the variants included in GWAS summary statistics. For simplicity, we show the mean numbers among three traits.

**Figure S6. Estimated prior parameters for genes across all tissues for IBD, SCZ and SBP.** The estimated prior parameters for genes across all 49 GTEx tissues for IBD, SCZ and SBP. Each dot represents the gene parameter for one of the tissues. "Proportion Causal" is the prior inclusion probability of genes. "Effect Size" is the prior variance for gene effects. "Enrichment" is the ratio of gene to variant prior inclusion probabilities. "PVE" is the proportion of variance explained by genes.

**Figure S7. Estimated proportions of mediated heritability by eQTLs for** SCZ (**A**) and SBP (**B**), as estimated by MESC (X-axis) and by cTWAS (Y-axis). Each dot represents the result from one GTEx tissue. The black dotted lines denote equivalence between the methods, and the blue dashed lines denote the slope relating cTWAS and MESC mediated heritability estimates. Top two tissues by cTWAS were highlighted.

**Figure S8. Number of genes detected by cTWAS, at PIP > 0.8, across all tissues,** for IBD (**A**), SCZ (**B**), and SBP (**C**). Each bar represents the result from one GTEx tissue.

**Figure S9. Comparison of number of genes detected by TWAS and cTWAS across all tissues for IBD (A), SCZ (B), and SBP (C).** cTWAS uses the cutoff of PIP > 0.8 (Y-axis), and TWAS uses a Bonferonni threshold (X-axis). Each dot represents the result from one GTEx tissue. The blue dashed lines denote the slopes relating the number of genes detected by cTWAS and TWAS.

**Figure S10. Number of genes detected by cTWAS, at PIP > 0.8, across tissue groups for SCZ (A) and SBP (B).** A gene was detected in a tissue group if it was detected in any of the tissues in that group. A gene was considered novel if it was not a previously reported candidate gene or if it was not the nearest gene to a genome-wide significant locus ("Nearest").

**Figure S11. Tissue specificity of genes detected by cTWAS.** For all genes detected at PIP > 0.8 in at least one GTEx tissue, the number of tissues where the genes were detected at a relaxed threshold of PIP > 0.5 were shown. (A) 110 genes for IBD. (B) 104 genes for SCZ. (C) 142 genes for SBP.

**Figure S12. cTWAS results at selected loci for IBD. A.** The *TYMP* locus using the "Esophagus Mucosa" eQTL data. The figure legend is the same as in the locus plots of the main figures (e.g. see Fig. 4B). **B.** The *LSP1* locus using the "Esophagus Muscularis" eQTL data. **C.** The *CCR5* locus using the "White Blood" eQTL data.

**Figure S13. Investigating false positives using PMR-Egger.** Detailed results for a single simulation from the high PVE scenario. **A.** Significance by TWAS and PMR-Egger for 6,454 genes analyzed by both methods. Vertical and horizontal dashed lines denote Bonferroni significance thresholds for TWAS and PMR-Egger, respectively. Diagonal dotted line denotes equivalence between the two methods. "Causal genes" have a true effect on the simulated trait. "Genes near causal variant(s)" are non-causal genes within 100kb of variants with true effect on the simulated trait. "Non-causal genes" are other genes not in these categories, including those potentially confounded by genes. **B.** GWAS and eQTL summary statistics for variants at the *ICA1* locus. The top panel shows the magnitude of the GWAS z-scores. The purple square is the TWAS z-score of the *ICA1* gene. The middle panel shows the magnitude of the weights from the FUSION lasso-based eQTL prediction model of *ICA1*. The bottom panel shows the magnitude of the eQTL z-scores for *ICA1*. In all panels, the solid red lines denote the location of a simulated causal variant. The black dashed lines denote the boundaries for PMR-Egger input (+/-100kb from gene body).

### Supplemental Tables

**Table S1. cTWAS results of all analyzed genes for LDL using liver eQTL data.** The "id" and "genename" columns are Ensembl gene IDs and gene symbols. The "chrom" and "pos" columns are the genomic positions of the first eQTL in the prediction model for each gene. Note that eQTL positions rather than TSS positions are reported here because cTWAS uses eQTL positions to assign genes to regions for analysis. The "region_tag" column is a unique identifier of the region that each gene was analyzed in.

The “cs_index” column denotes the confidence set that each gene is assigned to within a region; values of 0 indicate genes that were either unassigned or assigned to an impure confidence set (see Methods). The “PIP”, “tau2”, and “PVE” columns are the posterior inclusion probabilities, effect sizes, and proportions of variance explained using cTWAS for each gene. The “z” and “num_eqtl” columns are the TWAS z-scores and the number of eQTL in the gene prediction models after harmonization for each gene.

**Table S2. cTWAS results of silver standard and bystander genes for LDL using liver eQTL data.** The “annotation” column denotes whether each gene is in the silver standard (“known”) or bystander (“bystander”) gene lists. Column legends for other columns are the same as in Table S1. Silver standard genes without imputed expression are only listed with gene symbols and their number of eQTL (0). All other columns are shown as “NA”s. The “z”-score column shows the results from standard TWAS analysis with a genome-wide, Bonferroni-corrected significance threshold of $|z| > 4.56$.

**Table S3. GO enrichment analysis of the genes detected by cTWAS at PIP > 0.8 for LDL cholesterol using liver eQTL data.** The “Term” and “DB” columns are the GO terms and database names. The “Overlap” column is the number of detected genes divided by the total number of genes in each term. The “P.value” and “Adjusted.P.value” columns are the p-values for enrichment of detected genes in each term before and after multiple testing correction using B-H. The “Odds.Ratio” column is the ratio of the odds of detected genes being in a term relative to the odds of undetected genes being in a term. The “Combined.Score” column is a metric reported by Enrichr for ranking enriched terms; it is the product of the log p-value and the z-score of deviation from expected rank for each term. The “Genes” column lists the detected genes under each term.

**Table S4. GO enrichment analysis of 69 silver standard genes for LDL cholesterol.** Column legends are the same as in Table S2.

**Table S5. GO enrichment analysis using MAGMA for LDL cholesterol.** The “Category” and “GeneSet” columns are the database names and GO terms. The “N_genes” and “N_overlap” columns are the total number of genes and the number of genes identified from the GWAS by MAGMA in each term. The “p” and “adjP” columns are the p-values for enrichment of genes identified from the GWAS in each term before and after multiple testing correction using B-H. The “genes” column lists the genes identified from the GWAS by MAGMA in each term.

**Table S6. Estimated parameters and proportion of mediated heritability using cTWAS across all tissues for IBD, SCZ, and SBP.** The “trait” column is the abbreviated trait name for each analysis. The “tissue” column is the tissue used in each analysis. The “prior_g” and “prior_v” columns are the estimated prior inclusion probabilities for genes and variants in each analysis. The “prior_var_g” and “prior_var_v” columns are the estimated prior effect sizes for genes and variants in each analysis. The “enrich” column is the ratio of “prior_g” to “prior_v” and is a measure of the importance of gene inclusion relative to variant inclusion. The “pve_g” and “pve_v” columns are the proportions of trait variance attributable to genes and variants in each analysis. The “h2” column is the total heritability from all genetic variants and genes, computed as the sum of “pve_g” and “pve_v”. The “prop_h2_g” column is the proportion of total heritability mediated by genes, computed as “pve_g” / “h2”.

**Table S7. Proportion of mediated heritability using MESC across all tissues for IBD, SCZ, and SBP.** The “trait” column is the abbreviated trait name for each analysis. The “weight” column is the tissue used in each analysis. The “pve_g” and “pve_v” columns are the proportions of trait variance attributable to genes and variants in each analysis. The “h2” column is the total heritability, computed as the sum of “h2med” and “h2nonmed”. The “prop_h2_g” column is the proportion of total heritability mediated by genes, computed as “h2med”/”h2”. Note that expression scores for “Kidney_Cortex” tissue were not available from MESC.

**Table S8. cTWAS results of all analyzed genes across all tissues for IBD, SCZ, and SBP.** Results are stored separately for each combination of traits and tissues. Column legends for each set of results are the same as in Table S1.

**Table S9. IBD genes detected by cTWAS at PIP > 0.8 in the Blood/Immune and Digestive tissue groups**. The “genename” and “ensembl_gene_id” columns are gene symbols and Ensembl gene IDs. The “chromosome” and “start_position” columns are the genomic positions of the transcription start site for each gene. The “max_pip tissue” and “max_pip” columns denote the tissue with the highest PIP for each gene and its corresponding PIP. The “region_tag_tissue” column is a unique identifier of the region that each gene was analyzed in, for the tissue with the maximum PIP. The “z_tissue” and “num_eqtl_tissue” columns are the TWAS z-scores and the number of eQTL in the gene prediction models after harmonization for each gene, for the tissue with the maximum PIP. The “nearest_region_peak” column indicates if each gene is the nearest gene to the maximum GWAS signal in the region. The “distance_region_peak” column is the number of bases to the maximum GWAS signal in the region of each gene. The “which_nearest_region_peak” column lists the gene(s) nearest the maximum GWAS signal in the region of each gene; “-” denotes an unnamed gene. The “nearby” column indicates if each gene is within 500kb of a genome-wide significant locus. The “nearest” column indicates if each gene is the nearest gene to a genome-wide significant locus. The “distance” column is the number of bases to the nearest genome-wide significant locus, for genes that are “nearby” a genome-wide significant locus. The “which_nearest” column lists the gene nearest to a genome-wide significant locus, for genes that are “nearby” a genome-wide significant locus. The “known” column indicates genes that are on the silver standard gene list for IBD. The “GO_cTWAS” column lists the GO terms significantly enriched for cTWAS genes (Table S10) that are associated with each gene. The “GO_silver” column lists the GO terms significantly enriched for silver standard genes (Table S11) that are associated with each gene. The “GO_MAGMA” column lists the GO terms significantly enriched using MAGMA (Table S12) that are associated with each gene. For all three GO annotations, a maximum of 5 significant terms per gene are shown, ordered by odds ratios (cTWAS, silver standard) or p-values (MAGMA).

**Table S10. GO enrichment analysis using WebGestalt of genes detected by cTWAS at PIP > 0.8 for IBD using eQTLs in the Blood/Immune or Digestive tissue groups.** The “geneSet” and “description” columns are the GO terms and names. The “size” column is the total number of genes in each term. The “overlap” column is the number of detected genes in each term. The “expect” column is the expected number of detected genes in each term. The “enrichment” column is the ratio of the “overlap” and “expect” columns. The “pValue” and “FDR” columns are the p-values for enrichment of detected genes in each term before and after correcting for multiple comparisons. The “Genes” column lists the detected genes in each term.

**Table S11. GO enrichment analysis using Enrichr of genes detected by cTWAS at PIP > 0.8 for IBD using eQTLs in the Blood/Immune or Digestive tissue groups.** Column legends are the same as in Table S3.

**Table S12. GO enrichment analysis of silver standard genes for IBD.** Column legends are the same as in Table S3.

**Table S13. GO enrichment analysis using MAGMA for IBD.** Column legends are the same as in Table S5.
