## Supplementary figures and images for "Adjusting for genetic confounders in transcriptome-wide association studies leads to reliable detection of causal genes"

### Figure_S1.pdf

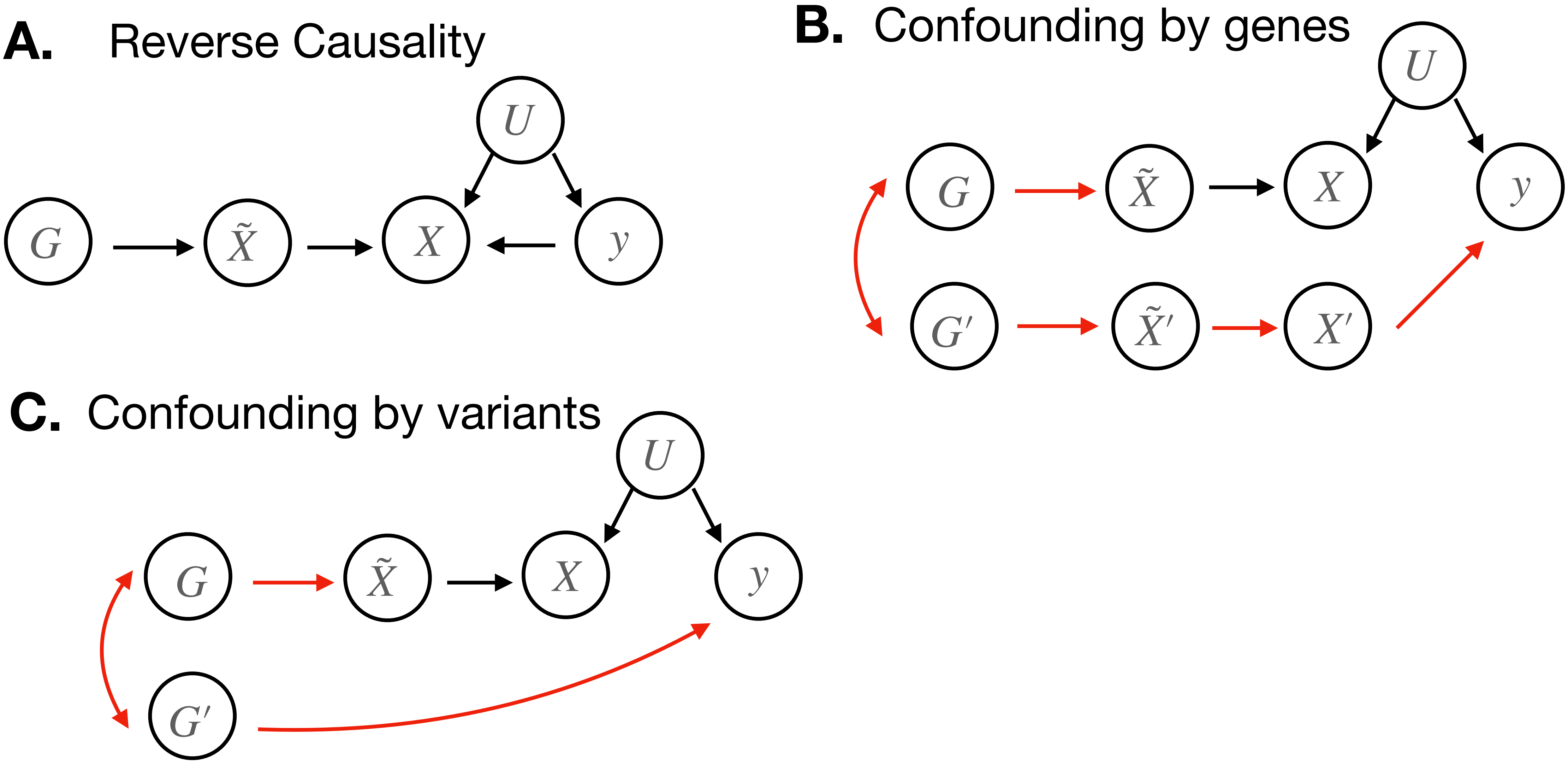

### Figure_S2.pdf

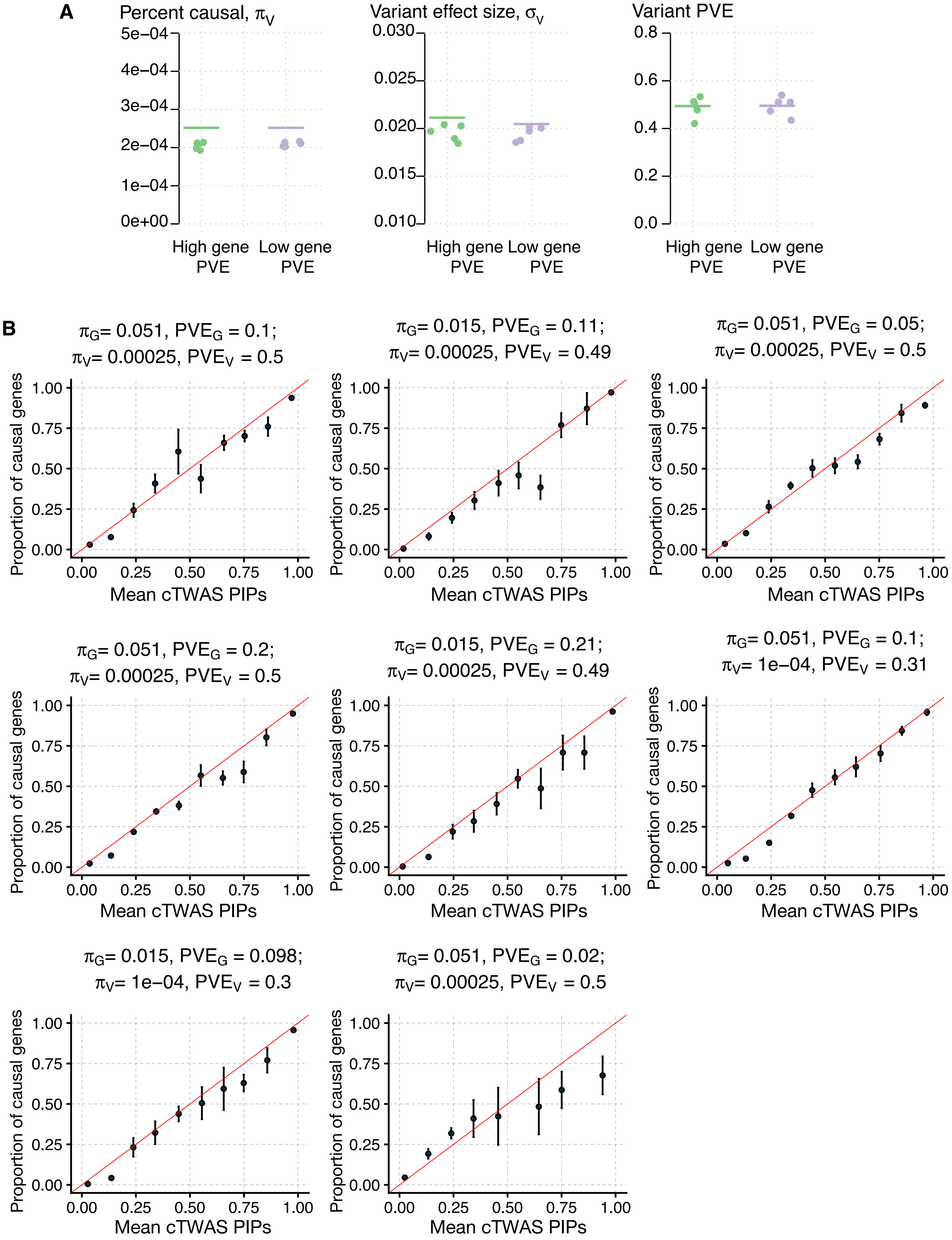

### Figure_S3.pdf

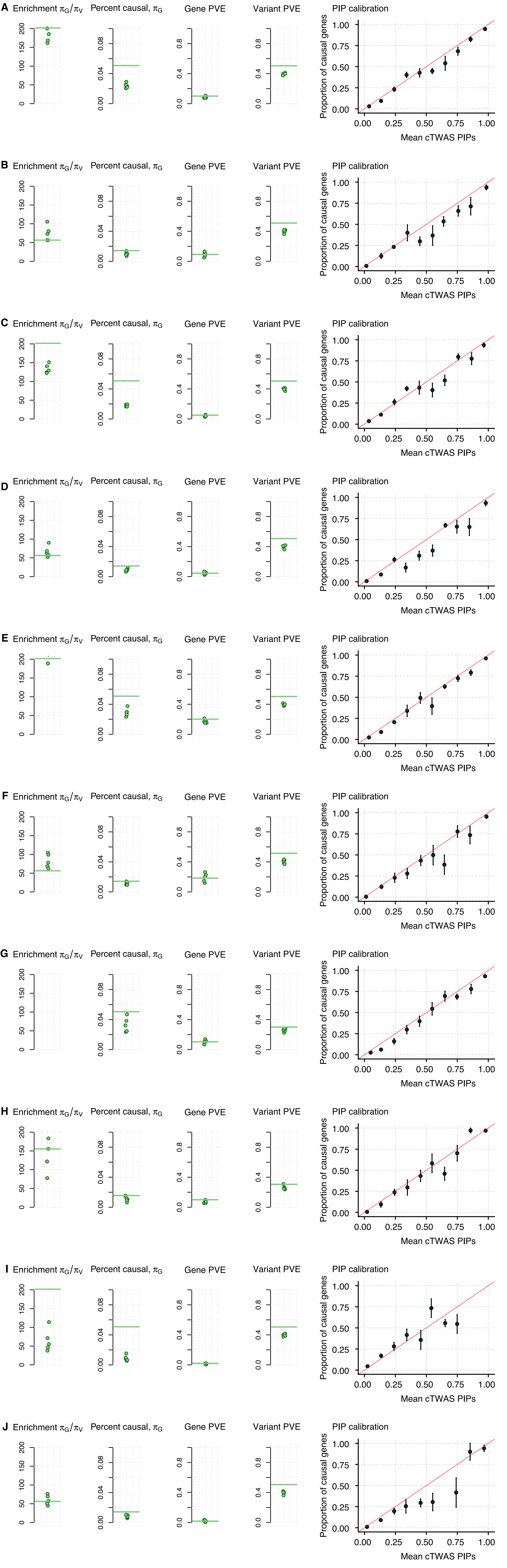

### Figure_S4.pdf

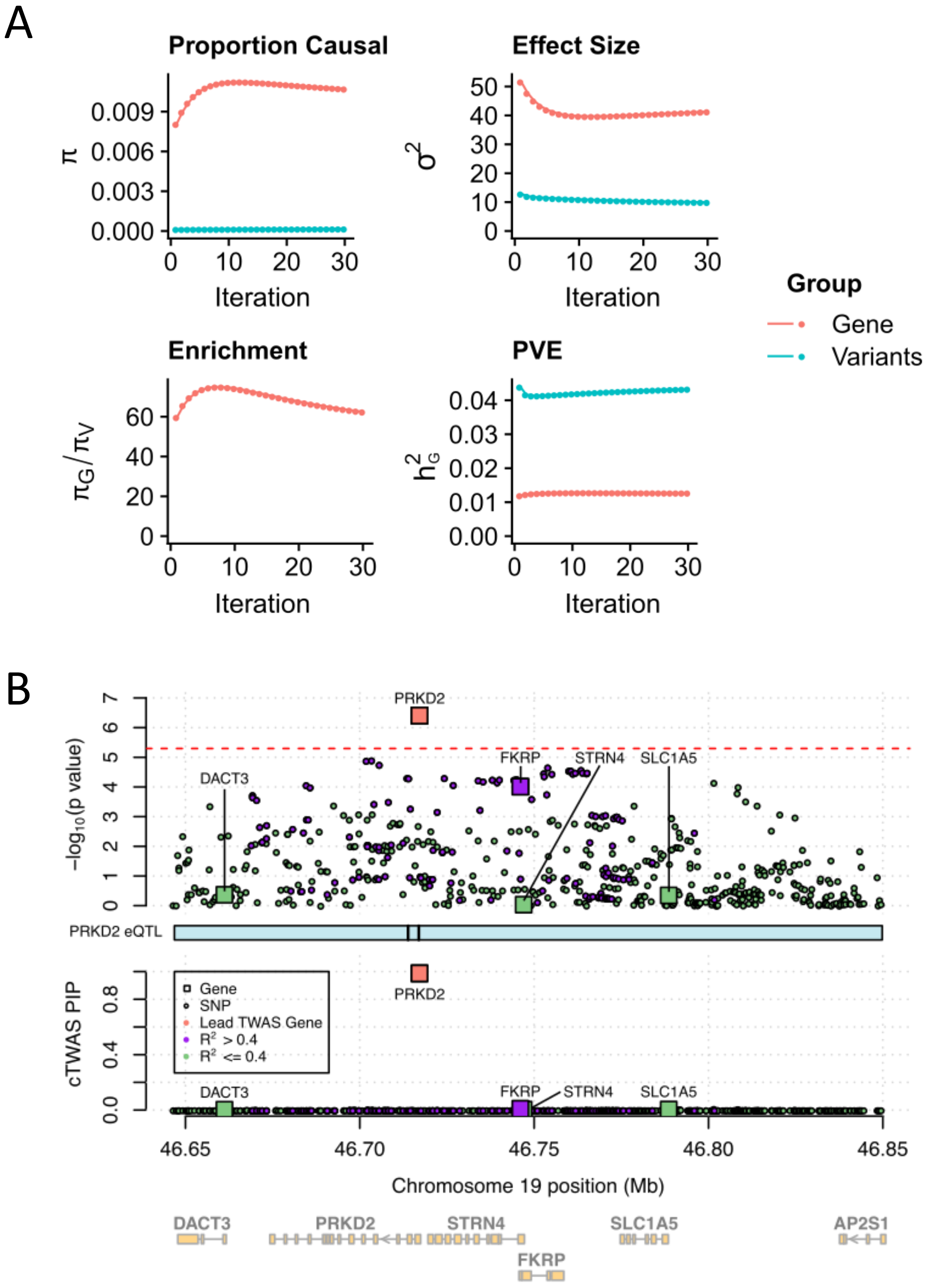

### Figure_S5.pdf

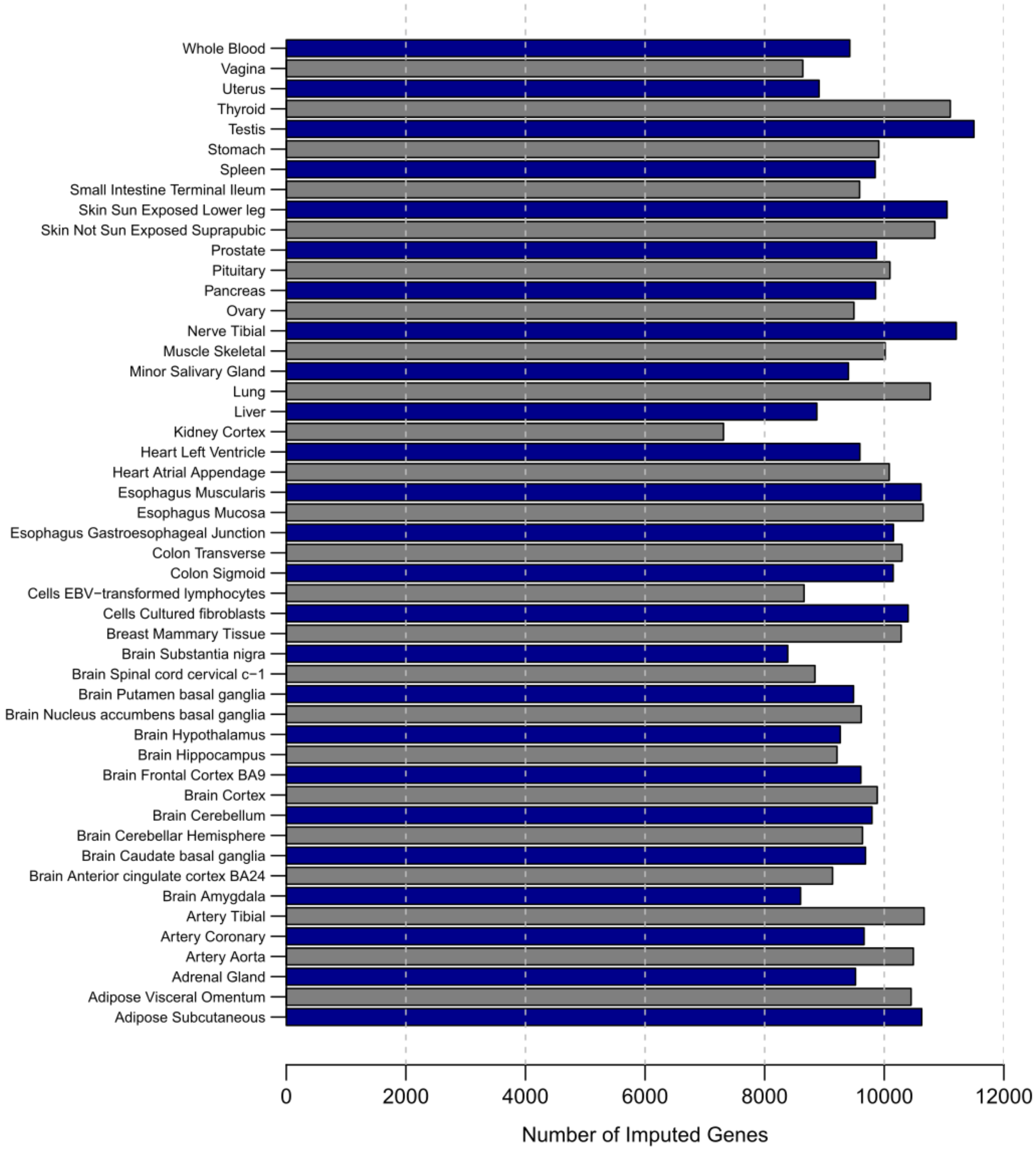

### Figure_S6.pdf

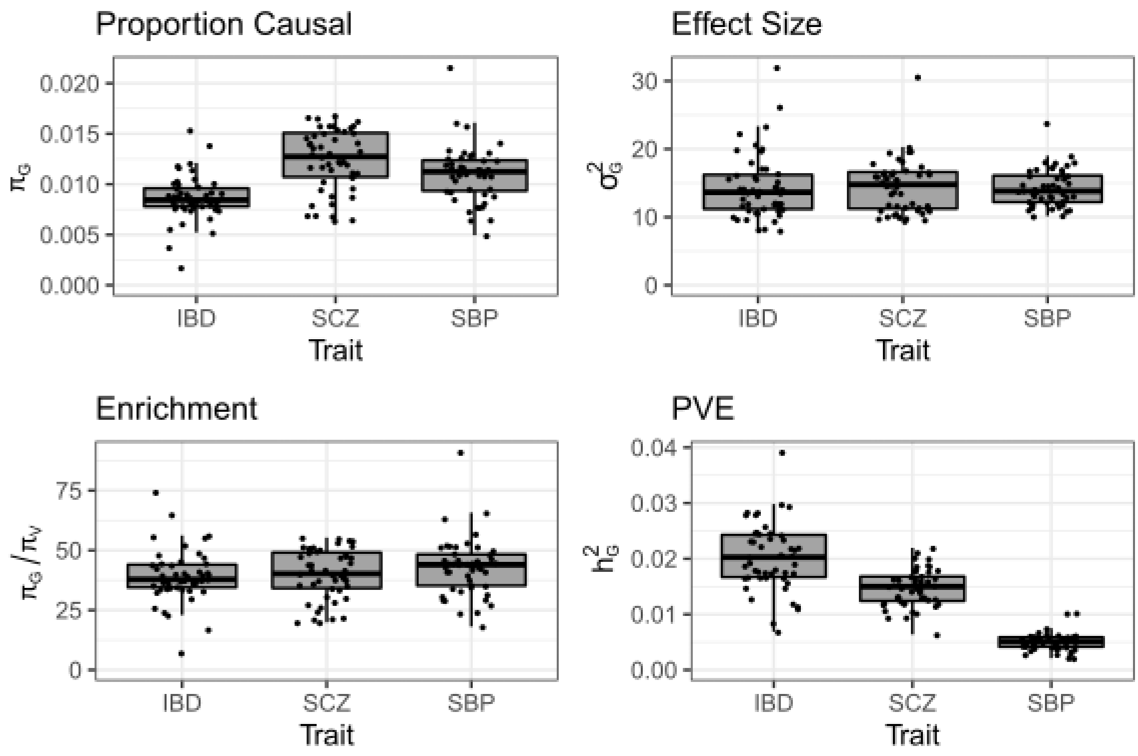

### Figure_S7.pdf

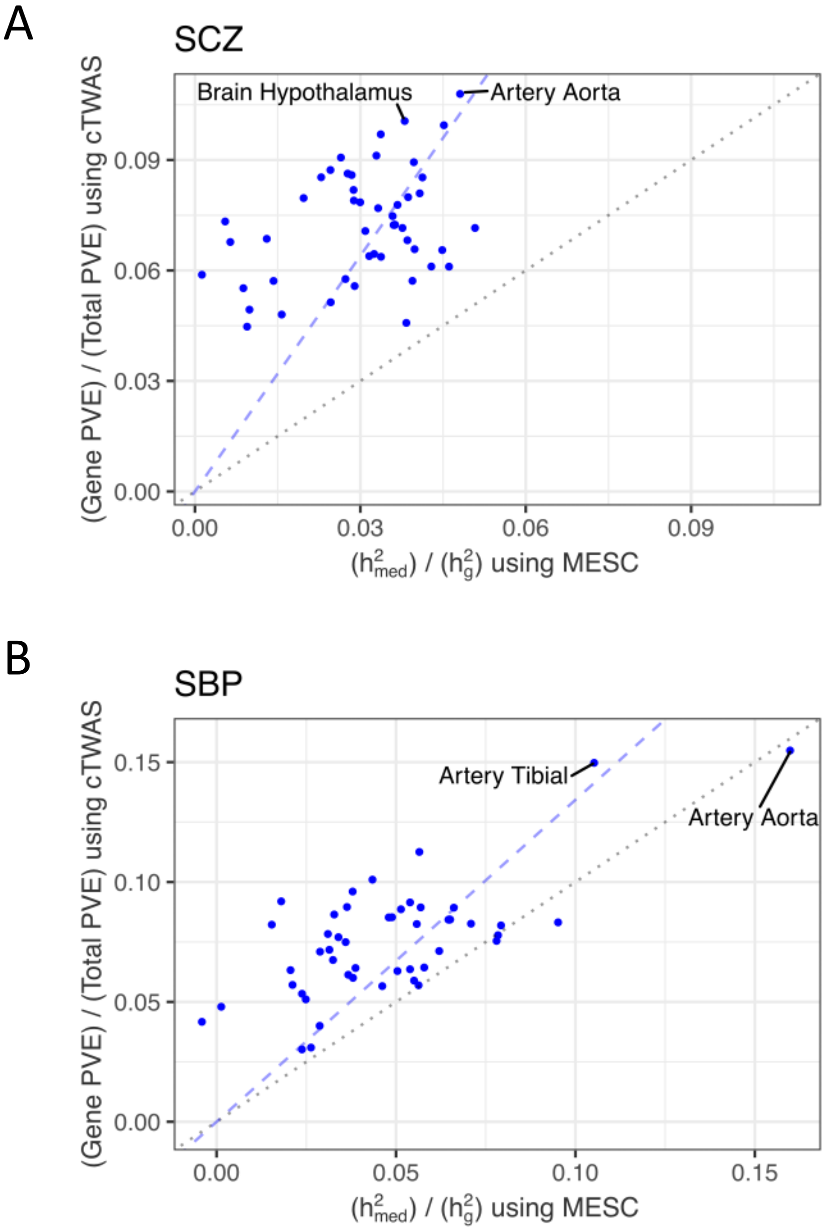

### Figure_S8a.pdf

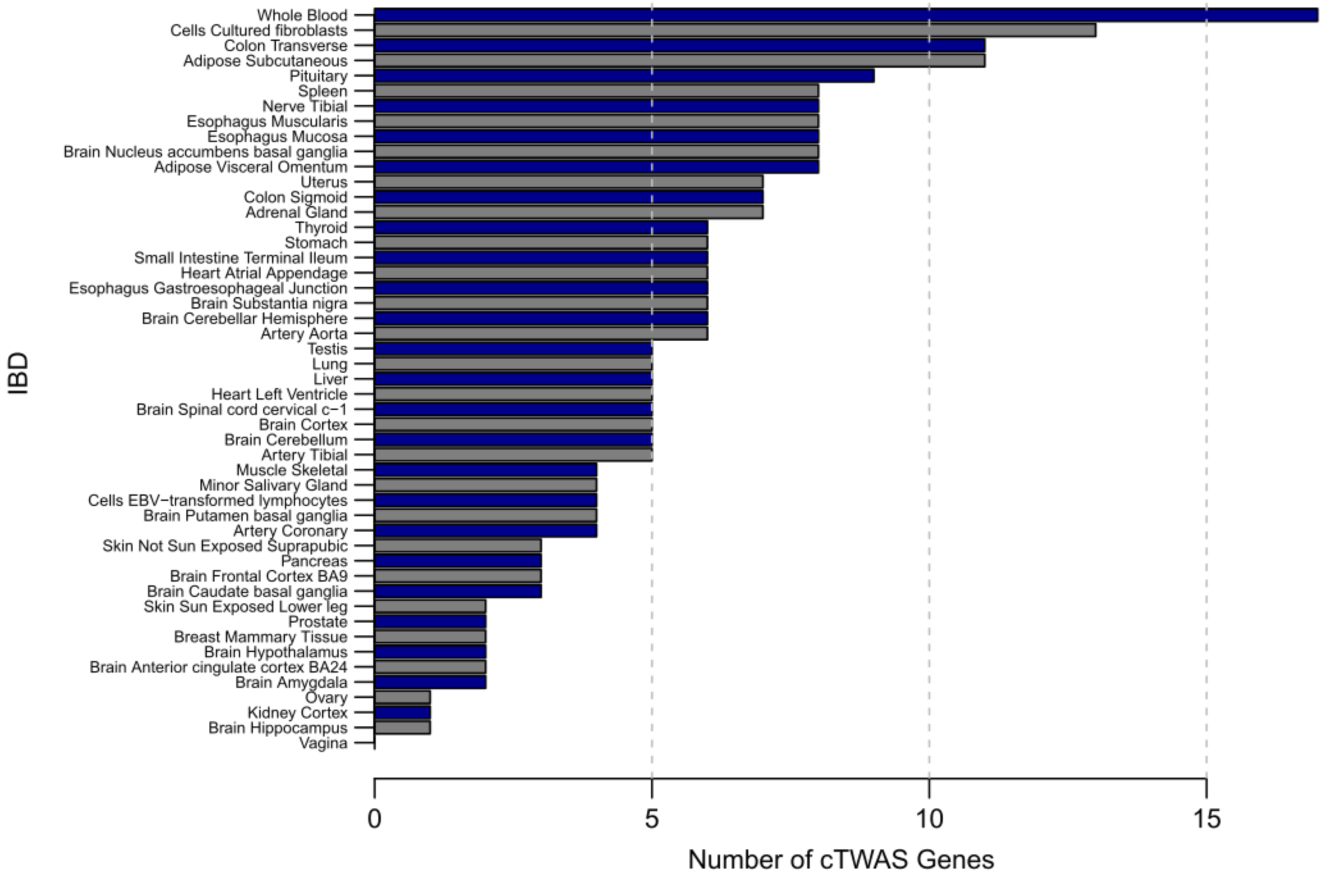

### Figure_S8b-c.pdf

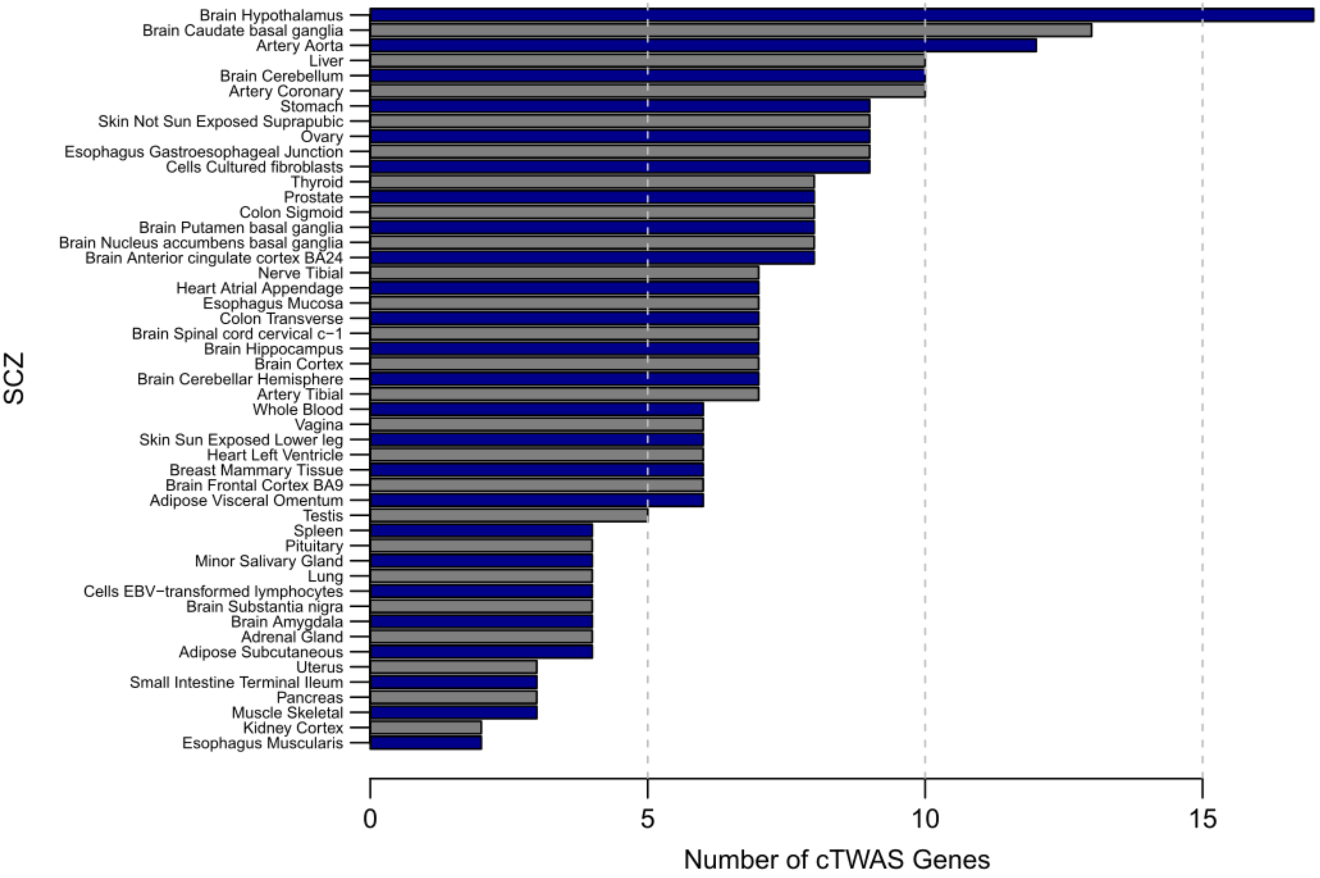

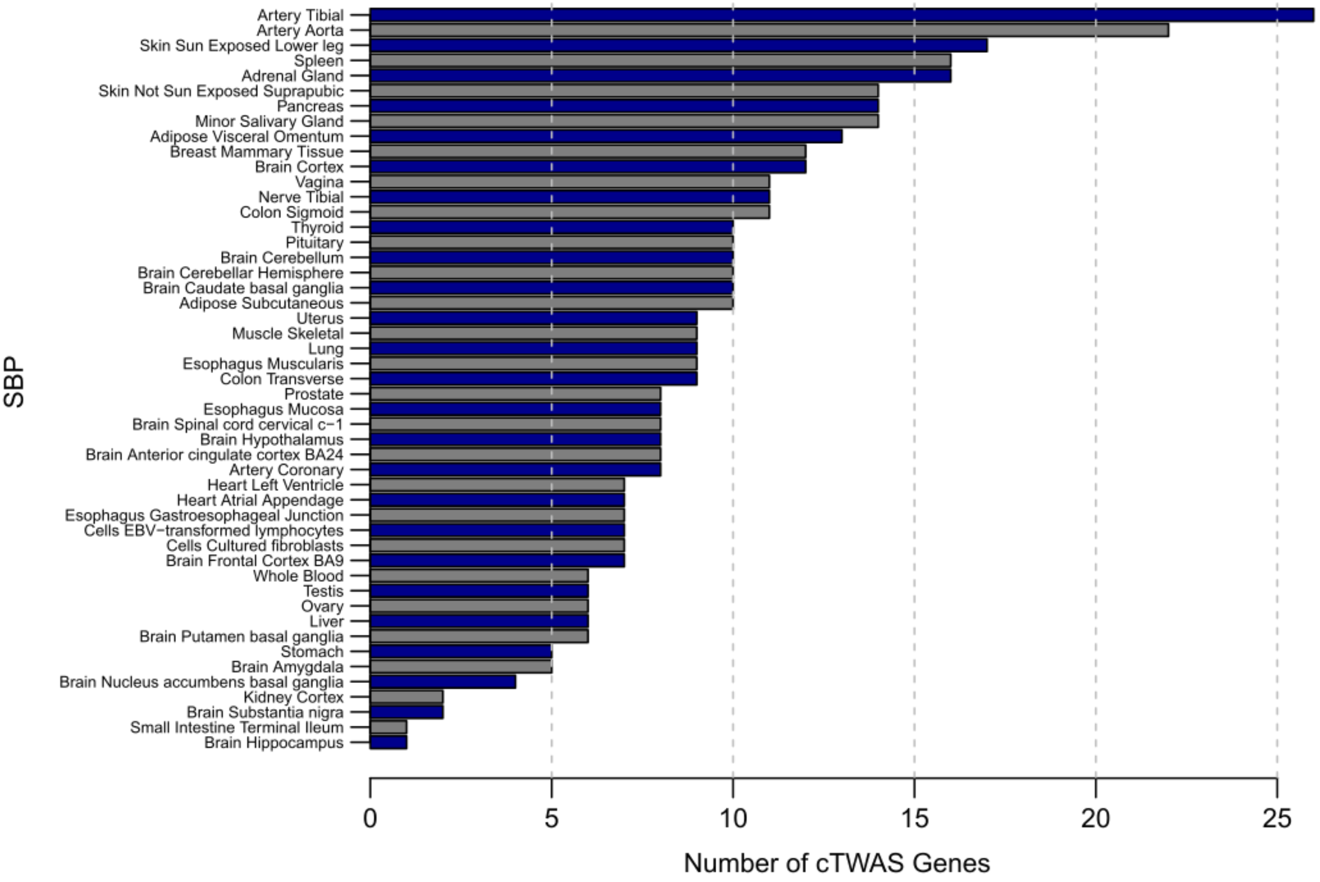

### Figure_S9.pdf

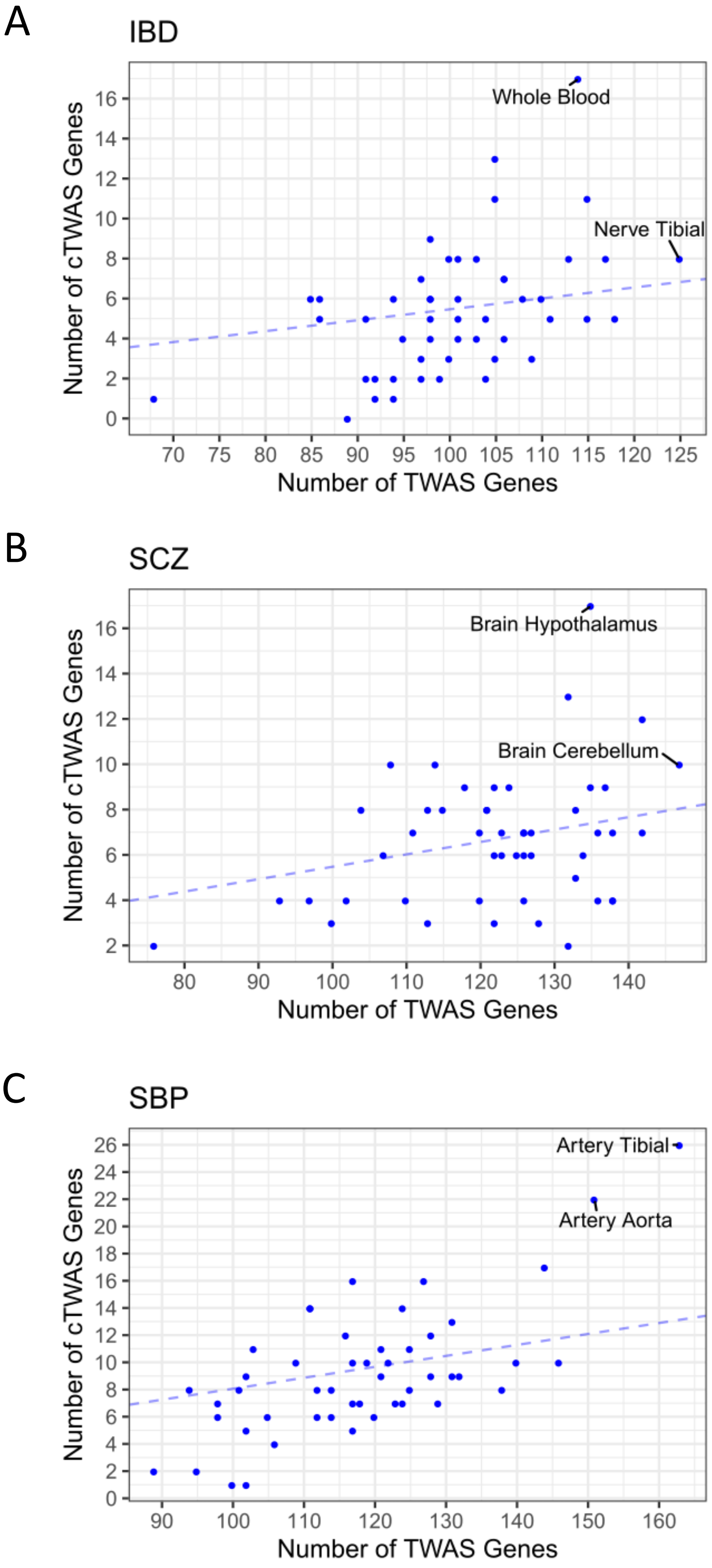

### Figure_S10.pdf

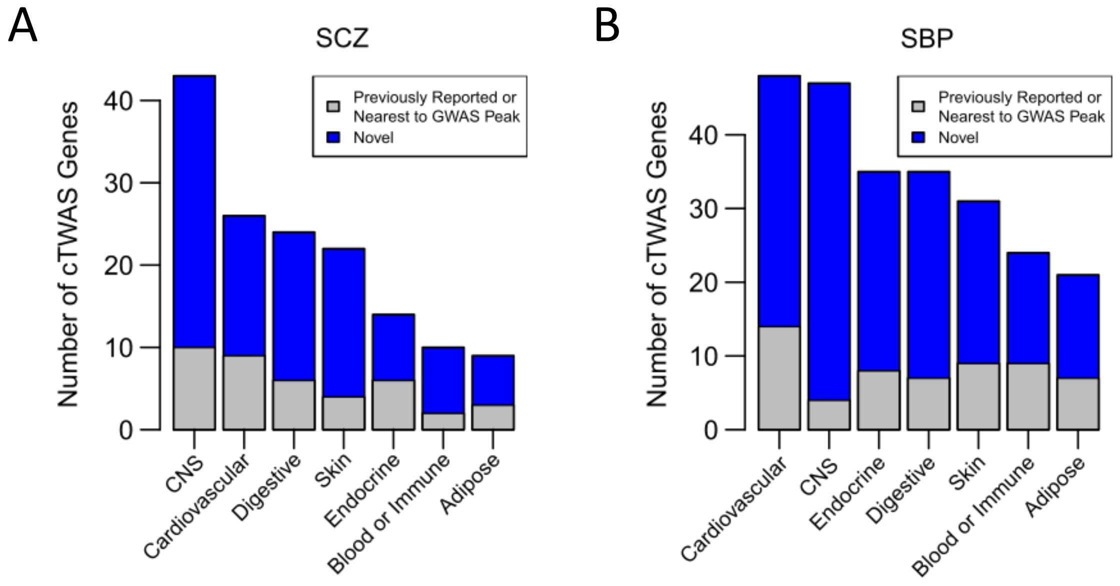

### Figure_S11.pdf

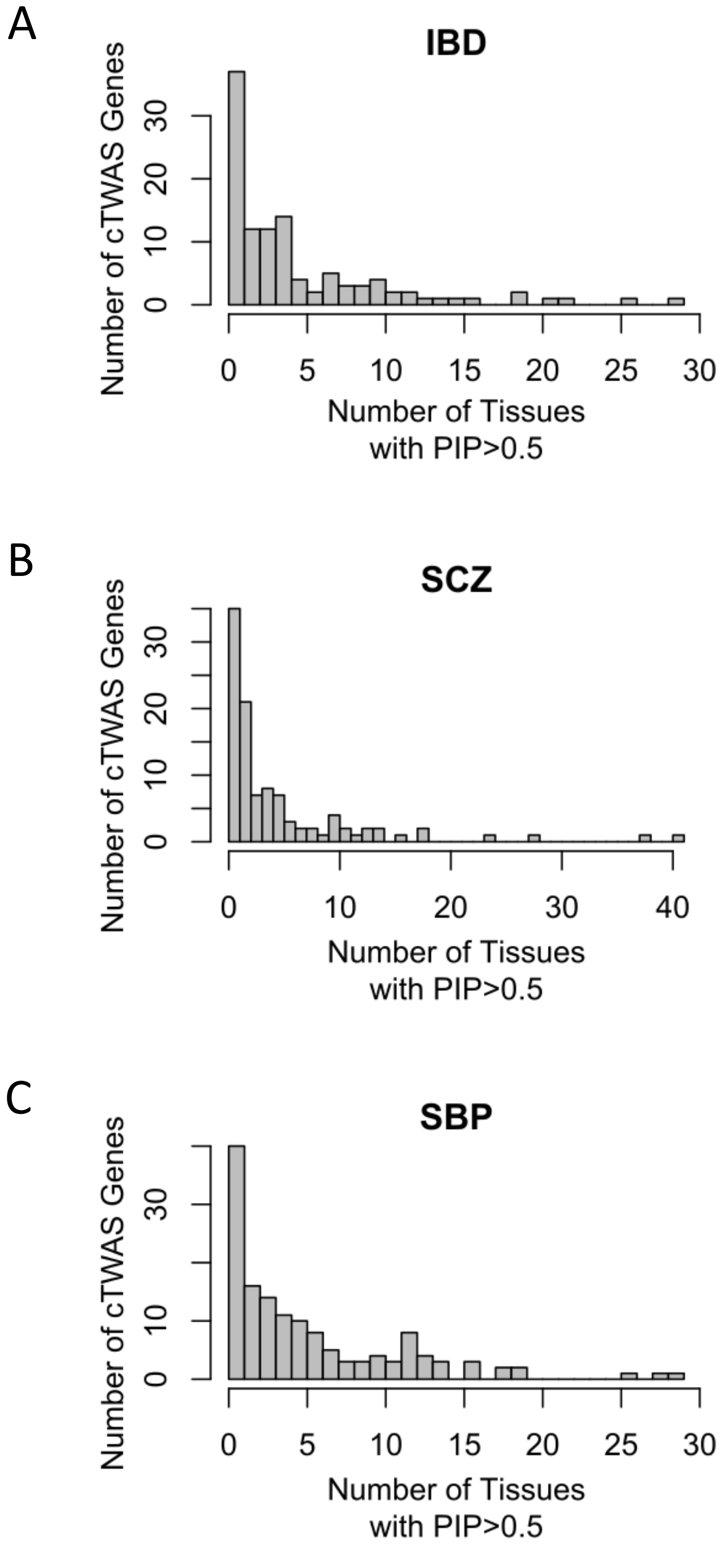

### Figure_S12.pdf

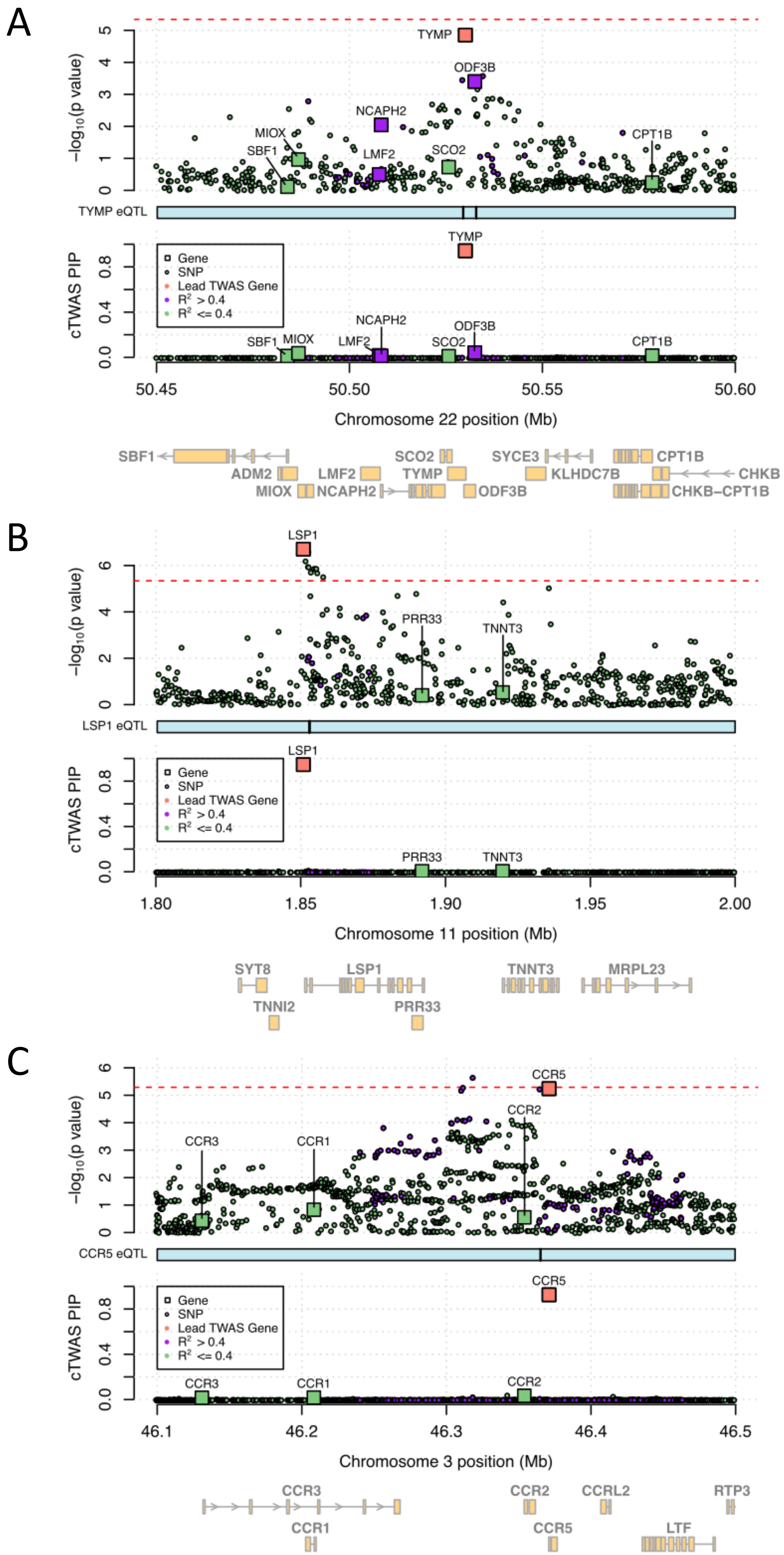

### Figure_S13.pdf

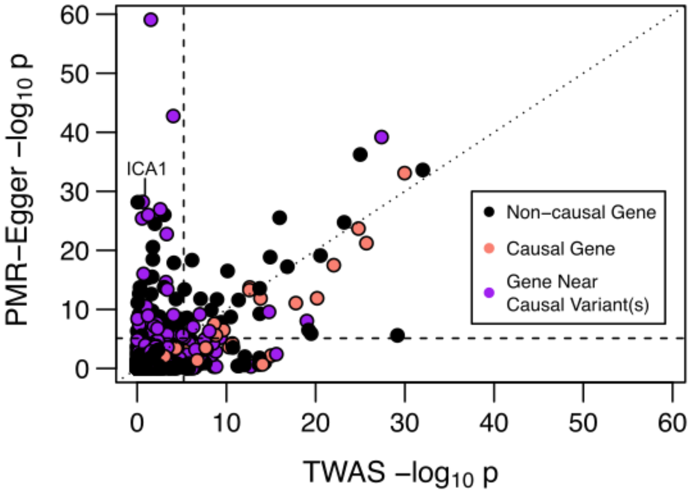

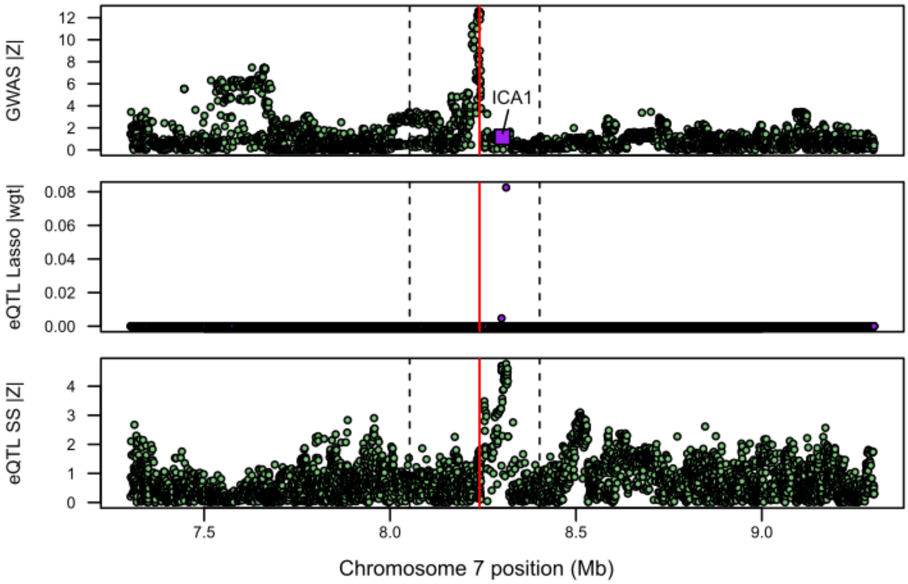
